# A modular platform for on-demand vaccine self-assembly enabled by decoration of bacterial outer membrane vesicles with biotinylated antigens

**DOI:** 10.1101/2021.08.24.457488

**Authors:** Kevin B. Weyant, Julie Liao, Mariela Rivera-De Jesus, Thapakorn Jaroentomeechai, Tyler D. Moeller, Steven Hoang-Phou, Sukumar Pal, Sean F. Gilmore, Riya Singh, David Putnam, Christopher Locher, Luis M. de la Maza, Matthew A. Coleman, Matthew P. DeLisa

**Affiliations:** Robert F. Smith School of Chemical and Biomolecular Engineering, Cornell University, Ithaca, NY 14853 USA; Nancy E. and Peter C. Meinig School of Biomedical Engineering, Cornell University, Ithaca, NY 14853 USA; Versatope Therapeutics, Inc., 110 Canal Street, Lowell, MA 01852 USA; Department of Pathology and Laboratory Medicine, Medical Sciences, Room D440, University of California, Irvine, Irvine, CA 92697 USA; Lawrence Livermore National Laboratory, Livermore, CA 94550 USA; Cornell Institute of Biotechnology, Cornell University, Ithaca, NY 14853 USA

## Abstract

Engineered outer membrane vesicles (OMVs) derived from laboratory strains of bacteria are a promising technology for the creation of non-infectious, nanoparticle vaccines against diverse pathogens. As mimics of the bacterial cell surface, OMVs offer a molecularly-defined architecture for programming repetitive, high-density display of heterologous antigens in conformations that elicit strong B and T cell immune responses. However, antigen display on the surface of OMVs can be difficult to control and highly variable due to bottlenecks in protein expression and localization to the outer membrane of the host cell, especially for bulky and/or complex antigens. To address this shortcoming, we created a universal approach called AddVax (avidin-based dock-and-display for vaccine antigen cross (x)-linking) whereby virtually any antigen that is amenable to biotinylation can be linked to the exterior of OMVs whose surfaces are remodeled with multiple copies of a synthetic antigen receptor (SNARE) comprised of an outer membrane scaffold protein fused to a member of the avidin family. We show that SNARE-OMVs can be readily decorated with a molecularly diverse array of biotinylated subunit antigens, including globular and membrane proteins, glycans and glycoconjugates, haptens, lipids, and short peptides. When the resulting OMV formulations were injected in wild-type BALB/c mice, strong antigen-specific antibody responses were observed that depended on the physical coupling between the antigen and SNARE-OMV delivery vehicle. Overall, these results demonstrate AddVax as a modular platform for rapid self-assembly of antigen-studded OMVs with the potential to accelerate vaccine generation, respond rapidly to pathogen threats in humans and animals, and simplify vaccine stockpiling.

## Introduction

Outer membrane vesicles (OMVs) are spherical bilayered nanostructures (~20-250 nm) ubiquitously released from the cell envelope of Gram-negative and Gram-positive bacteria and their production represents a *bona fide* bacterial secretion process (*1, 2*). As derivatives of the cell envelope, OMVs mimic the structural organization and conformation of the bacterial cell surface while also containing periplasmic lumenal components. Natively produced OMVs mediate diverse functions such as increasing pathogenicity in the host environment (*3*), promoting bacterial survival under conditions of stress (*4*), and controlling interactions within microbial communities (*5*).

In addition to their natural biological roles, OMVs have enabled a spectrum of bioengineering applications, most notably in drug and vaccine delivery, that exploit the unique structural and functional attributes of these nanoparticle systems (*6–9*). OMVs are especially attractive as a vaccine platform because they are non-replicating, immunogenic facsimiles of the producing bacteria and thus contain the pathogen-associated molecular patterns (PAMPs) present on bacterial outer membranes (*6, 7*). These PAMPs endow OMVs with intrinsic immunostimulatory properties that strongly stimulate innate and adaptive immune responses (*10–13*). In addition to this in-built adjuvanticity, OMVs are right-sized for direct drainage into lymph nodes and subsequent uptake by antigen presenting cells and cross-presentation (*14*). From a translational perspective, OMVs can be readily produced at high quantities and commercial scales via standard bacterial fermentation, and their clinical use has already been established in the context of OMVs from pathogenic *Neisseria meningitidis* serogroup B (MenB), also known as outer membrane protein complexes (OMPCs), that are the basis of a polyribosylribitol phosphate (PRP) conjugate vaccine approved for *Haemophilus influenzae* type b called PedvaxHIB^®^ (*15*) and are a component of the MenB vaccine Bexsero^®^ (*16*).

To generalize and expand the vaccine potential of OMVs, we and others have leveraged recombinant DNA technology and synthetic biology techniques to engineer OMVs with heterologous protein and peptide cargo (*17, 18*). By targeting expression to the outer membrane or the periplasm of an OMV-producing host strain, both surface display as well as payload encapsulation are possible, providing versatility as biomedical research tools and vaccines. Typically, this involves genetic fusion of a protein or peptide of interest (POI) to an outer membrane scaffold protein (e.g., the *E. coli* cytolysin ClyA), with the resulting POI accumulating in released OMVs that can be readily recovered from the culture supernatant. These methods have made it possible to enlist non-pathogenic, genetically tractable bacteria such as *Escherichia coli* K-12 for the production of designer OMVs that are loaded with foreign antigens of interest (*6, 19*). When inoculated in mice, such engineered OMVs stimulate antigen-specific humoral B cell and dendritic cell (DC)-mediated T cell responses including activation of CD4^+^ and CD8^+^ T cells (*10, 20-22*). Importantly, the immune responses triggered by antigen-loaded OMV vaccines have proven to be protective against a range of foreign pathogens including bacteria and viruses (*23–26*) as well as against malignant tumors (*27*). While proteins and peptides remain the focus of most OMV-based vaccine efforts, advances in bacterial glycoengineering have enabled decoration of OMV exteriors with heterologous polysaccharide antigens, giving rise to a new class of glycoconjugate vaccines that can effectively deliver pathogen-mimetic glycan epitopes to the immune system and confer protection to subsequent pathogen challenge (*28–30*). Collectively, these and other studies have revealed that the repetitive, high-density arrangement of antigens on the OMV surface enhances the response to otherwise poorly immunogenic epitopes such as small peptides and polysaccharides, which likely results from induction of strong B-cell receptor clustering.

These successes notwithstanding, the classical approach to loading OMVs with foreign antigens prior to their isolation from bacterial cultures is not without its challenges. For example, many antigens that are desirable from a vaccine standpoint are incompatible with recombinant expression in the lumen or on the surface of OMVs. While there can be many reasons for this, the most common bottlenecks include misfolding, proteolytic degradation, and/or inefficient bilayer translocation of the POI, especially for those that are very bulky and/or structurally complex. Because there are currently no effective tools for predicting *a priori* the expressibility of OMV-directed antigens, the creation of heterologous OMV vaccines remains very much a time-consuming trial-and-error process that often must be repeated for each new antigen. Even when a foreign antigen can be successfully localized to OMVs, it may lack important post-translational modifications that are formed inefficiently (or not at all) in the bacterial expression host. In addition, it can be difficult or even impossible to precisely control the quantity of OMV-associated antigen, thereby excluding antigen density as a customizable design parameter. It should also be noted that while it is possible to integrate polypeptide and polysaccharide biosynthesis with the vesiculation process (*6, 19*), it has yet to be demonstrated whether biosynthesis of other biomolecules can be similarly integrated, thereby limiting the spectrum of cargo that can be packaged in OMVs.

To address these shortcomings, reliable strategies are needed for modular OMV functionalization in which OMV vectors and structurally diverse target antigens are separately produced and then subsequently linked together in a controllable fashion. Along these lines, direct chemical conjugation of proteins and polysaccharides to OMVs/OMPCs following their isolation has been reported (*15, 31*); however, this technique involves non-specific attachment of antigens to unknown OMV components and thus is heterogeneous and difficult to predict or analyze. Moreover, non-uniform coupling of antigen to particulate carriers may result in sub-optimal immunogenicity. For more precise, homogenous antigen attachment, site-specific conjugation methods are preferrable. To this end, two groups recently demonstrated specific bioconjugation on OMVs by adapting a “plug-and-display” approach that had previously been developed for decorating virus-like particles with protein and peptide antigens (*32*). This approach involved the use of the SpyTag/SpyCatcher protein ligation system to covalently attach purified SpyTag-antigen (or SpyCatcher-antigen) fusion proteins onto cognate SpyCatcher-scaffold (or SpyTag-scaffold) fusions that were expressed on the surface of OMVs (*33, 34*). While this enabled loading of exogenous antigens on OMVs, with one report even demonstrating specific anti-tumor immune responses (*33*), the protein ligation strategy is limited to proteinaceous antigens that are compatible with isopeptide bond formation.

To develop a more universal strategy for tethering virtually any biomolecular cargo to the exterior of OMVs, we created AddVax (avidin-based dock-and-display for vaccine antigen cross (x)-linking) whereby biotinylated antigens are linked to the exterior of ready-made OMVs whose surfaces are remodeled with biotin-binding proteins. The method involves producing OMV vectors that repetitively display multiple copies of a synthetic antigen receptor (SNARE) comprised of an outer membrane scaffold protein fused to a member of the avidin family. Following their production and isolation, SNARE-OMVs can be readily decorated with a wide range of biotinylated subunit antigens, including globular and membrane proteins, glycans and glycoconjugates, haptens, lipids, and short peptides. Importantly, antigen-studded SNARE-OMVs promote strong antigen-specific antibody responses that compare favorably to the responses measured for classically prepared OMV formulations (i.e., cellular expression of antigen-scaffold fusions). Overall, our results demonstrate that AddVax is a highly modular and versatile platform for on-demand vaccine creation that should enable rapid cycles of development, testing, and production of new OMV-based vaccines for numerous diseases.

## Results

### A modular framework for self-assembly of antigens on OMV surfaces

As a first step towards developing a universal platform for rapidly assembling antigens of interest on the surface of OMVs, we constructed SNAREs by translationally fusing a cell surface scaffold protein to a biotin-binding protein (**Fig. 1a**). A panel of cell surface scaffold modules were chosen based on their ability to direct passenger proteins to the *E. coli* outer membrane. These included cytolysin ClyA (*18*), hybrid protein Lpp-OmpA (*35*), and the autotransporter β-domains derived from the N-terminus of intimin (Int) (*36*) and the C-termini of adhesin involved in diffuse adherence (AIDA-I), antigen-43 (Ag43), hemoglobin-binding protease (Hbp), and immunoglobulin A protease (IgAP) (*37*). Initially, each scaffold was fused in-frame to enhanced monoavidin (eMA) (**Fig. 1b**), a derivative of dimeric rhizavidin (RA) that was designed to be monomeric with highly stable, biotin-binding properties (*38*), and subsequently expressed from the arabinose-inducible plasmid pBAD24 in hypervesiculating *E. coli* strain KPM404 Δ*nlpI* (*39*). This strain is an endotoxin-free BL21(DE3) derivative (sold as ClearColi™ by Lucigen) that we previously engineered to vesiculate through knockout of the *nlpI* gene (*40*). Using this strain, OMVs were readily produced that contained full-length SNARE chimeras, with Lpp-OmpA-eMA and Int-eMA showing the strongest expression albeit with significant amounts of higher and lower molecular weight species that likel corresponded to aggregation and degradation products, respectively (**Supplementar Fig. 1a**).

**Figure 1.**
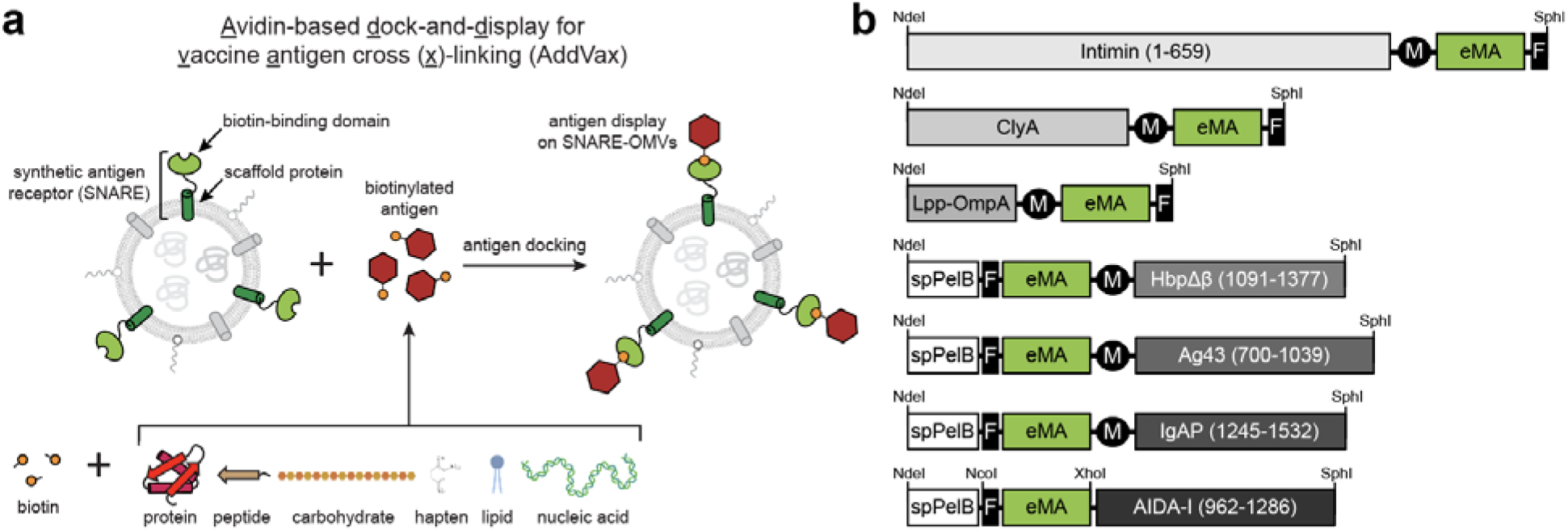
A modular platform for rapid self-assembly of OMV-based vaccine candidates. (a) Schematic of AddVax technology whereby ready-made OMVs displaying a synthetic antigen receptor (SNARE-OMVs) are remodeled with biotinylated antigens-of-interest. Using AddVax, the surface of SNARE-OMVs can be remodeled with virtually any biomolecule that is amenable to biotinylation including peptides, proteins, carbohydrates, glycolipids, glycoproteins, haptens, lipids, and nucleic acids. (b) Genetic architecture of SNARE constructs tested in this study. Numbers in parentheses denote amino acids of the scaffold that were fused to the biotin-binding eMA domain and used for membrane anchoring. Additional features include: export signal peptide from PelB (spPelB); c-Myc epitope tag (M); FLAG epitope tag (F), and NdeI, SphI, and NcoI restriction enzyme sites used for cloning.

To evaluate antigen docking, we initially focused on biotinylated green fluorescent protein (biotin-GFP) as the target antigen (**Supplementary Fig. 2a**), which enabled facile and quantitative prototyping of the different SNARE-OMV designs. When biotin-GFP was incubated with 100 ng SNARE-OMVs coated on ELISA plates, all exhibited dose-dependent binding up to ~10 nM of biotin-GFP except for the eMA-AIDA-Iβ and ClyA-eMA receptors, which appeared saturated at low levels of biotin-GFP (**Supplementar Fig. 1b**). The lack of binding for these two SNAREs was not entirely surprising given that these constructs exhibited very weak expression compared to the other SNAREs (**Supplementar Fig. 1b**). Importantly, there was no detectable binding of unmodified GFP by any of the SNARE-OMVs, indicating that antigen capture was entirely dependent upon the presence of the biotin moiety. Next, the two most effective SNAREs in terms of biotin-GFP binding, namely eMA-IgAPβ and Lpp-OmpA-eMA, were evaluated over a range of conditions to identify parameters (e.g., growth temperature, culture density at time of induction, inducer level, etc.) that affected GFP docking levels (**Supplementary Results**and **Supplementar Fig. 3**). Overall, the engineered Lpp-OmpA-eMA receptor outperformed eMA-IgAPβ in terms of biotin-GFP binding capacity (**Supplementar Fig. 3a**); however, expression of this construct from pBAD24 using standard amounts of L-arabinose (0.2% or 13.3 mM) was detrimental to the host cells based on the observation that the final culture densities hardly changed, and in some cases even decreased, from the densities at the time of induction, which was not the case for eMA-IgAPβ (**Supplementar Fig. 3b**). Given the different biogenesis pathways of the IgAP autotransporter versus the Lpp-OmpA β-barrel outer membrane protein, we suspected that the host cell toxicity associated with Lpp-OmpA might result from inducer levels that were too strong. In support of this notion, when Lpp-OmpA-eMA constructs were induced with ~50-times less inducer, the post-induction cell growth was markedly improved, with Lpp-OmpA-eMA-expressing cells reaching a final density on par with that of cells expressing eMA-IgAPβ (**Supplementary Fig. 3c**). Importantly, the Lpp-OmpA-eMA SNARE-OMVs isolated from these healthier host cells captured significantly more biotin-GFP compared to eMA-IgAPβ SNARE-OMVs. An even higher level of biotin-GFP binding was obtained by moving the Lpp-OmpA-eMA construct into the L-rhamnose-inducible plasmid pTrham (**Supplementary Fig. 3d-f**), which is known for its tighter expression control compared to pBAD vectors and can help to overcome the deleterious saturation of membrane and secretory protein biogenesis pathways (*41, 42*). To determine the effect of the biotin-binding module on antigen loading and to further highlight the modularity of AddVax, we constructed a panel of Lpp-OmpA-based SNAREs comprised of alternative biotin-binding domains including dimeric RA, tetrameric streptavidin (SA), and monomeric streptavidin with a lowered off-rate (mSA^S25H^) (*43*). The SNAREs comprised of RA and mSA^S25H^ both captured biotin-GFP at a level that was nearly identical to the eMA-based receptor, while the SA-based receptor showed binding that was barely above background, a result that appears to be due to the poor expression of this SNARE compared to the others (**Supplementar Fig. 4a and b**). Given the similarity in antigen capture efficiency for the eMA, RA, and mSA^S25H^ SNAREs as well as post-induction culture growth (**Supplementary Fig. 4a**), we chose the more extensively characterized Lpp-OmpA-eMA SNARE (expressed from plasmid pTrham with 0.5 mM L-rhamnose inducer) for all further studies.

### Antigen loading capacity of SNARE-OMVs

To determine the loading capacity of Lpp-OmpA-eMA SNARE-OMVs, the OMV fractions were first subjected to extensive washing with ultracentrifugation to recover washed OMVs, and then irreversible aggregates were removed by filtration through 0.45 μm pores. Next, we quantified bound antigen by mixing Lpp-OmpA-eMA SNARE-OMVs with biotin-GFP in solution and subsequently measuring the amount of OMV-bound GFP in an ELISA-style format. This assay was designed to mirror the process of vaccine self-assembly, whereby ready-made SNARE-OMVs are mixed with biotinylated antigens in an on-demand fashion. Importantly, the dose-response profile for pre-binding biotin-GFP on SNARE-OMVs in solution was in relative agreement with the dose-response curve generated by capturing biotin-GFP on the surface of immobilized SNARE-OMVs (**Fig. 2a**). The maximum amount of biotin-GFP that was captured on the SNARE-OMV surface was ~1% by mass when ~2 wt % biotin-GFP was input to the mixture, with the addition of higher amounts of biotin-GFP leading to no significant increase in biotin-GFP binding (**Fig. 2b**). In both assay formats, the combination of SNARE-OMVs with non-biotinylated GFP or biotin-GFP with blank OMVs lacking a SNARE resulted in little to no detectable binding (**Fig. 2a, b**). We also found that the maximum biotin-GFP loading on SNARE-OMVs was lower but on par with the amount of GFP that was displayed on the surface of OMVs following cellular expression of a scaffold-antigen fusion, ClyA-GFP (**Fig. 2c**) (*18*). Despite this difference, an advantage of SNARE-OMVs was the ability to vary the antigen loading density over a wide biotin-GFP concentration range, thereby providing a level of control that is more difficult to achieve with cellular expression of scaffold-antigen fusions. When visualized by transmission electron microscopy (TEM), SNARE-OMVs decorated with biotin-GFP had a size (~50 nm) and overall appearance that was indistinguishable from unloaded SNARE-OMVs (**Fig. 2d**) and consistent with previous TEM images of engineered OMVs (*18, 22, 25*) including those from the same hypervesiculating host strain used here (*40*). These findings indicate that controllable vesicle loading could be achieved using the AddVax approach without significantly impacting OMV ultrastructure.

**Figure 2.**
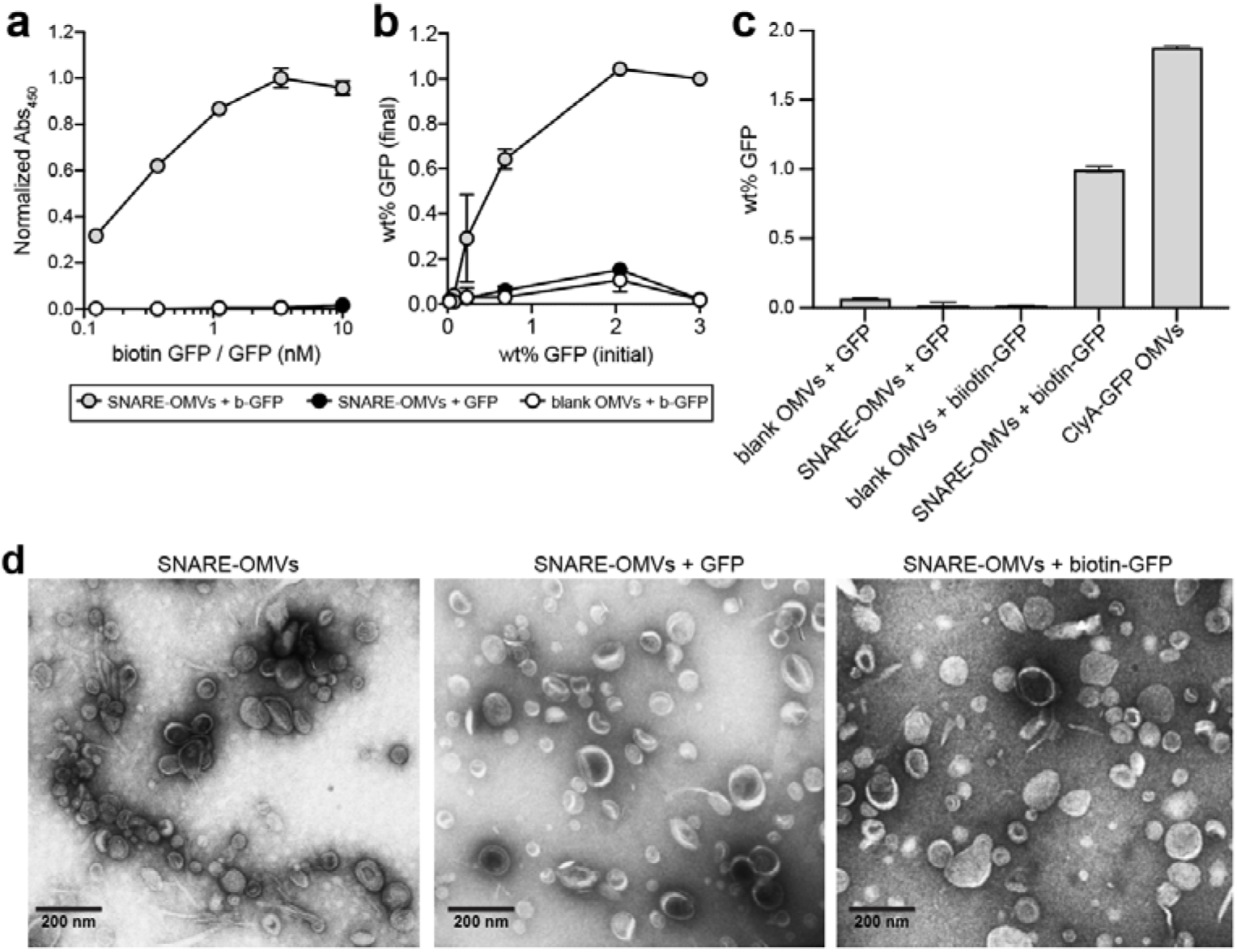
Chimeric Lpp-OmpA-eMA SNARE enables controllable antigen loading on OMVs. (a) Dose-response curve generated by loading biotin-GFP or unmodified GFP on SNARE-OMVs isolated from hypervesiculating *E. coli* strain KPM404 Δ*nlpI* expressing the Lpp-OmpA-eMA construct from plasmid pTrham (induced with 0.5 mM L-rhamnose). Blank OMVs were isolated from plasmid-free KPM404 Δ*nlpI* cells. Binding activity was determined by ELISA in which Lpp-OmpA-eMA SNARE-OMV were immobilized on plates and subjected to varying amounts of biotin-GFP, after which plates were extensively washed prior to detection of bound biotin-GFP using anti-polyhistidine antibody to detect C-terminal 6xHis tag on GFP. Data were normalized to the maximum binding signal corresponding to Lpp-OmpA-eMA SNARE-OMVs in the presence of 3.3 nM biotin-GFP. (b) Same OMVs as in (a) but dose-response was generated by first incubating OMVs with biotin-GFP or unmodified GFP in solution, washing to remove unbound protein, and determining GFP levels by ELISA-based detection. (c) Comparison of GFP levels on Lpp-OmpA-eMA SNARE-OMVs versus ClyA-GFP OMVs. ClyA-GFP OMVs were isolated from KPM404 Δ*nlpI* cells expressing ClyA-GFP fusion construct from plasmid pBAD18 as described in Kim et al. (*18*). Binding data are the average of triplicate measurements, and all error bars represent the standard deviation of the mean. (d) Transmission electron micrograph of Lpp-OmpA-eMA SNARE-OMVs alone or following incubation with unmodified GFP or biotin-GFP as indicated. The scale bar represents 200 nm.

### Decoration of OMV surfaces with structurally diverse antigens

To demonstrate the universality of the approach, we next investigated decoration of SNARE-OMVs with a diverse array of biotinylated antigens. Some of these were chosen because their incorporation into the OMV structure through cellular expression as a scaffold-ant fusion protein was predicted to be difficult or impossible. For example, *Plasmodium falciparum* Pfs25 protein (Pfs25), a glycophosphotidylinositol (GPI)-anchored protein expressed on the surface of zygotes and ookinetes, is a promising malaria transmission-blocking vaccine antigen (*44*). However, Pfs25 could not be expressed in soluble form in *E. coli* likely due to its 11 disulfide bonds (*45*), and thus is incompatible with conventional cellular expression techniques for OMV engineering. Along similar lines, *Chlamydia* major outer membrane protein (MOMP) is a β-barrel integral membrane protein (IMP) that accounts for ~60% of the mass of the outer membrane of *Chlamydia* spp. (*46, 47*) and is highly antigenic (*48*), making it an attractive subunit vaccine candidate (*49*). However, expression of MOMP in the *E. coli* cytoplasm results in aggregation and the formation of inclusion bodies (*50, 51*) while expression in the *E. coli* outer membrane results in significant cell toxicity (*50, 52, 53*). To incorporate these two challenging membrane protein antigens into SNARE-OMVs required generation of soluble versions of each antigen. For Pfs25, soluble expression was achieved using a baculovirus-insect cell expression system (**Supplementary Fig. 2b**), while for MOMP from *Chlamydia trachomatis* mouse pneumonitis (MoPn) biovar (strain Nigg II; now called *Chlamydia muridarum*), soluble expression was achieved using a protein engineering technology known as SIMPLEx (solubilization of IMPs with high levels of expression) (*54*) in which sandwich fusion between an N-terminal “decoy” protein, namely *E. coli* maltose-binding protein (MBP), and C-terminal truncated human apolipoprotein AI (ApoAI*) transformed *C. muridarum* MOMP (Cm-MOMP) into a water-soluble protein that was expressed at high levels in the *E. coli* cytoplasm (**Supplementar Fig. 2a**). Following incubation of SNARE-OMVs with biotinylated versions of insect cell-derived Pfs25 and *E. coli*-derived SIMPLEx-Cm-MOMP (Sx-Cm-MOMP), we observed efficient OMV decoration that depended on both the presence of the chimeric Lpp-OmpA-eMA receptor on OMVs and the biotin moiety on each antigen (**Fig. 3a and b**). In the case of Sx-Cm-MOMP, we observed a low but reproducible signal for both controls (SNARE-OMVs with non-biotinylated Sx-Cm-MOMP and blank OMVs with biotinylated Sx-Cm-MOMP) that may correspond to a small amount of auto-insertion of Sx-Cm-MOMP into OMVs.

**Figure 3.**
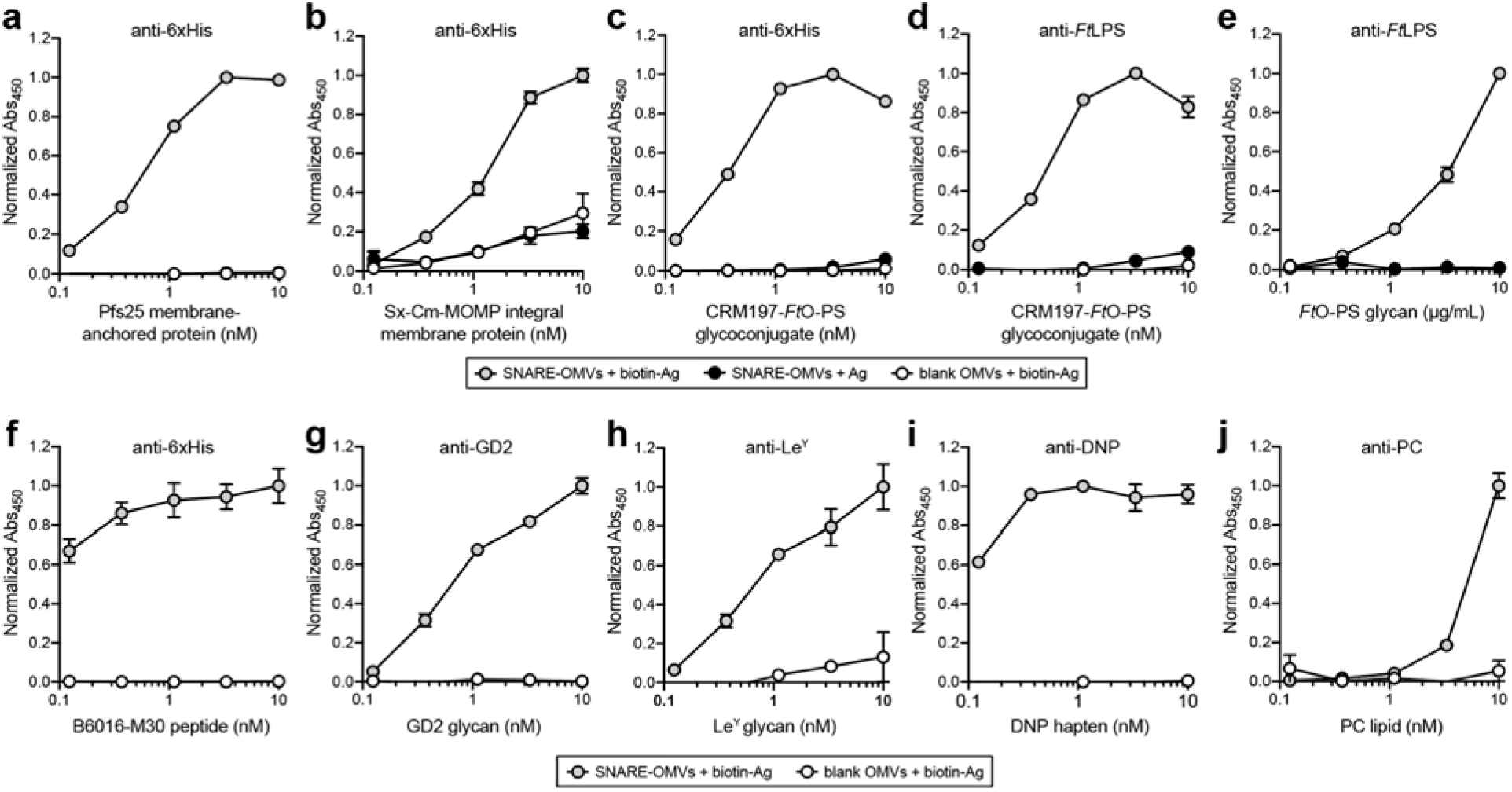
Rapid self-assembly of OMV vaccine candidates decorated with diverse biomolecular antigens. (a-j) Dose-response curves generated by loading biotinylated or non-biotinylated antigens on SNARE-OMVs isolated from hypervesiculating KPM404 Δ*nlpI* cells expressing the Lpp-OmpA-eMA construct from plasmid pTrham (induced with 0.5 mM L-rhamnose). Blank OMVs were isolated from plasmid-free KPM404 Δ*nlpI* cells. Binding activity was determined by ELISA in which Lpp-OmpA-eMA SNARE-OMVs were immobilized on plates and subjected to varying amounts of unbiotinylated or biotinylated antigen, after which plates were extensively washed prior to detection of bound antigen using the antibodies indicated at top of each panel. Data were normalized to the maximum binding signal in each experiment. All binding data are the average of triplicate measurements and error bars represent the standard deviation of the mean.

We next investigated carbohydrate structures such as lipopolysaccharide (LPS) antigens that represent appealing molecules for vaccine development owing to their ubiquitous presence on the surface of diverse pathogens and malignant cells. A challenge faced with most polysaccharides is that they make poor vaccines on their own because they are unable to interact with the receptors on T cells in germinal centers (GCs) (*55*). This can be overcome by covalent attachment of a polysaccharide to a carrier protein, which provides T cell epitopes that can induce polysaccharide-specific IgM-to-IgG class switching, initiate the process of affinity maturation, and establish long-lived memory (*56*). Despite the widespread success of glycoconjugates, there is an unmet need to identify formulations that elicit stronger primary antibody responses after a single immunization, especially in primed or pre-exposed adolescents and adults, and achieve prolonged vaccine efficiency (*56*). To this end, we speculated that AddVax would provide a convenient strategy for combining glycoconjugates with the intrinsic adjuvant properties of OMVs (*10–12*). Such an approach would provide a simpler alternative than attempting to combine OMV biogenesis with cellular expression of glycoconjugate vaccine candidates in *E. coli* (*57*), a feat that has yet to be reported. Thus, we attempted to adorn SNARE-OMVs with biotinylated glycoconjugates by leveraging an engineered *E. coli* strain (*58*) to produce the carrier protein CRM197 that was glycosylated at its C-terminus with a recombinant mimic of the *Francisella tularensis* SchuS4 O-antigen polysaccharide (*Ft*O-PS) (**Supplementary Fig. 2c**). Decoration of SNARE-OMVs with a biotinylated version of this glycoconjugate was readily detected by immunoblotting against both the CRM197 carrier and its covalently linked *Ft*O-PS antigen (**Fig. 3c and d**, respectively). We also demonstrated an alternative strategy for combining OMVs with polysaccharide antigens whereby a biotinylated version of *F. tularensis* SchuS4 LPS (*Ft*LPS) was directly bound to the exterior of SNARE-OMVs (**Fig. 3e**). This formulation was motivated by the fact that a protein providing T cell help only needs to be in close proximity to the polysaccharide in order to target the same B cell and does not have to be covalently linked to the polysaccharide to induce class switching and T-cell activation (*28, 29, 59*). Indeed, the co-delivery of non-covalently linked proteins and polysaccharides present on the exterior of OMVs is sufficient to make a polysaccharide immunogenic (*15, 28, 29*).

The final group of antigens that we investigated were small-sized biomolecules that are known to be weakly immunogenic by themselves and therefore require carrier molecules to increase chemical stability and adjuvanticity for the induction of a robust immune response. This group included: (i) B16-M30 peptide, a CD4^+^ T cell neoepitope expressed in the B16F10 melanoma as a consequence of a mutation in the *kif18b* gene (*60*); (ii) ganglioside GD2 glycan, a pentasaccharide antigen found on human tumors including melanoma, neuroblastoma, osteosarcoma, and small-cell lung cancer, that was highly ranked (12 out of 75) in a National Cancer Institute pilot program that prioritized the most important cancer antigens (*61*); (iii) Lewis Y (Le^Y^), a tetrasaccharide extension of the H blood group galactose-glucosamine that has been shown to be overexpressed on tumors (*62*); (iv) 2,4-dinitrophenol (DNP), a model hapten to which the immune system is unresponsive (*63*); and (v) phosphocholine (PC), a major lipid component of myelin and one of the main antigenic targets of the autoimmune response in multiple sclerosis, with lipid-reactive antibodies likely contributing to disease pathogenesis (*64*). In each case, we observed clearly detectable antigen binding on the surface of SNARE-OMVs that was significantly above the background seen with blank OMVs lacking biotin-binding receptors (**Fig. 3f-j**). Collectively, these results illustrate the potential of the AddVax approach for modular self-assembly of candidate OMV vaccines decorated with diverse biomolecular cargo.

### Immunogenicity of SNARE-OMVs loaded with model GFP antigen

We next sought to assess the immunological potential of SNARE-OMVs displaying biotin-GFP. Specifically, BALB/c mice were immunized via subcutaneous (s.c.) injection of SNARE-OMVs decorated with biotin-GFP or other control formulations after which blood was collected at regular intervals. Negative controls included blank SNARE-OMVs, SNARE-OMVs that were mixed with non-biotinylated GFP, and PBS. ClyA-GFP-containing OMVs generated by cellular expression, which were previously reported to elicit high antibody titers following immunization in mice (*22*), served as a positive control. Importantly, SNARE-OMVs displaying biotin-GFP elicited robust IgG responses to GFP that were significantly higher (*p* < 0.0001) than the titers measured for control mice immunized with blank SNARE-OMVs or PBS (**Fig. 4a**). It is particularly noteworthy that the total IgG titers triggered by SNARE-OMVs were indistinguishable from those measured in response to ClyA-GFP-containing OMVs, validating the antigen docking strategy as a potent alternative to cellular expression of scaffold-antigen fusions for boosting the immunogenicity of foreign subunit antigens, in particular those that are weakly immunogenic on their own such as GFP (*22, 65*). Interestingly, the IgG response elicited by non-tethered GFP that was mixed with SNARE-OMVs gave a significantly lower (*p* < 0.01) antigen-specific IgG response compared to biotin-GFP that was docked on OMVs, indicating that the physical coupling of the antigen to the surface of the OMV is essential for exploiting the full intrinsic adjuvanticity of OMVs. To determine whether the immune responses were Th1 or Th2 biased (*66*), IgG antibody titers were further broken down by analyzing IgG1 and IgG2a subclasses. Mice immunized with different GFP-containing OMVs showed robust mean titers of both GFP-specific IgG1 and IgG2a antibodies (**Fig. 4b**). For the groups immunized with ClyA-GFP OMVs, the relative titers of IgG1 and IgG2a subclasses were comparable, consistent with our earlier work and in line with responses typically seen with traditional subunit vaccines (*21*). In contrast, biotin-GFP-studded SNARE-OMVs elicited an IgG2a-dominant humoral response, suggesting induction of a Th1-biased immune response consistent with heightene cellular immunity stimulation.

**Figure 4.**
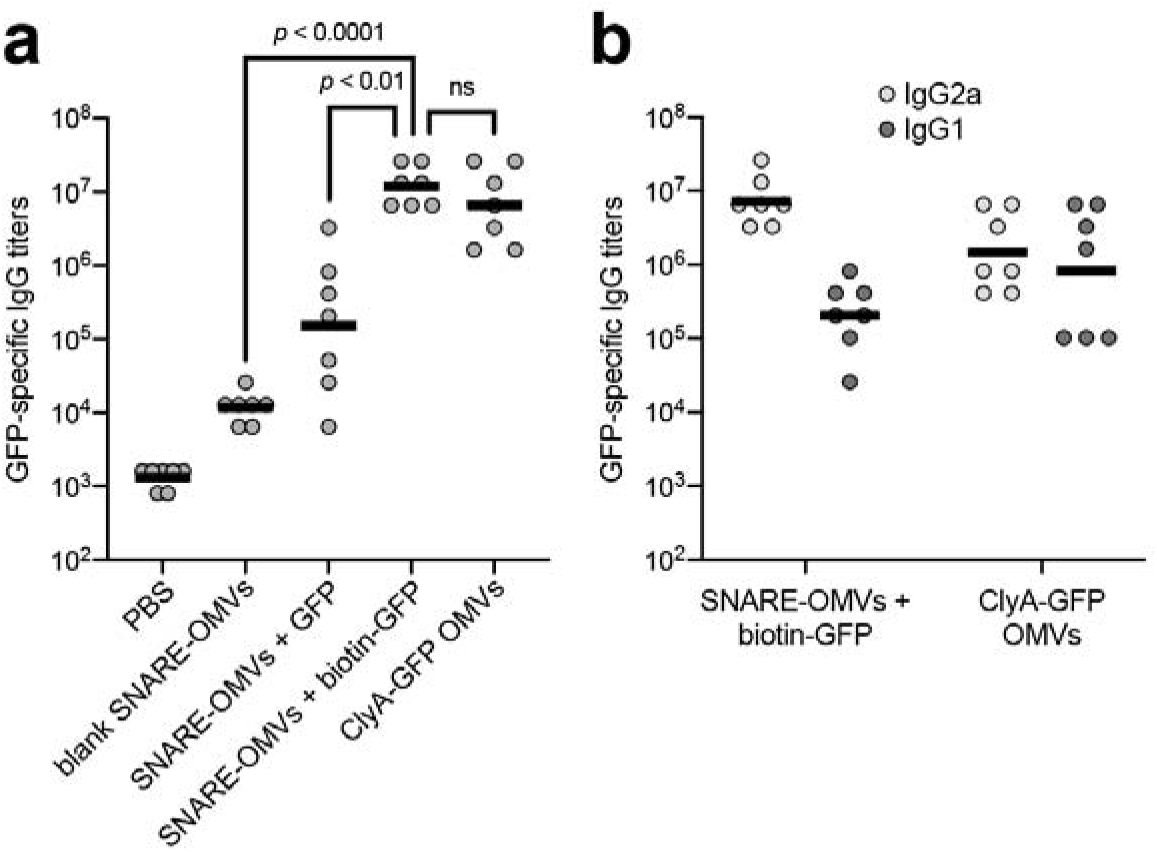
SNARE-OMVs decorated with biotin-GFP boost GFP-specific IgG titers. (a) GFP-specifi IgG titers in endpoint (day 56) serum of individual mice (gray dots) and geometric mean titers of each group (horizontal black lines). Five groups of BALB/c mice, seven mice per group, immunized s.c. with the following: PBS, SNARE-OMVs isolated from KPM404 Δ*nlpI* cells expressing the Lpp-OmpA-eMA construct, SNARE-OMVs mixed with non-biotinylated or biotinylated GFP, and ClyA-GFP isolated from KPM404 Δ*nlpI* cells expressing ClyA-GFP fusion. Mice received prime injections containing an equivalent amount of OMVs (20 μg total protein) on day 0 and were boosted on day 21 and 42 with the same doses.(b) Geometric mean IgG subclass titers measured from endpoint serum with IgG1 titers in dark gray and IgG2a in light gray. Statistical significance of antibody titers for SNARE-OMVs + biotin-GFP against blank SNARE-OMVs and SNARE-OMVs + GFP indicates statistically significant difference (*p* < 0.0001 and *p* < 0.01, respectively; unpaired *t* test with Welch’s correction) between the groups; ns – not significant.

### Immunogenicity of SNARE-OMVs loaded with validated Cm-MOMP antigen

Encouraged by the immunostimulation observed for SNARE-OMVs remodeled with the model GFP antigen, we next investigated the humoral immune response to SNARE-OMVs that were decorated with Cm-MOMP, a validated subunit vaccine candidate (*49, 51*). Prior to immunization, we first tested the antigenicity of our Sx-Cm-MO construct (**Fig. 5a**) that was engineered as described above for soluble, high-level expression. Immunoblots of purified Sx-Cm-MOMP were probed with anti-Cm-MOMP-specific monoclonal antibody (mAb) MoPn-40, which was generated by inoculation of BALB/c mice with *C. muridarum* followed by isolation of hybridomas producin antibodies against

**Figure 5.**
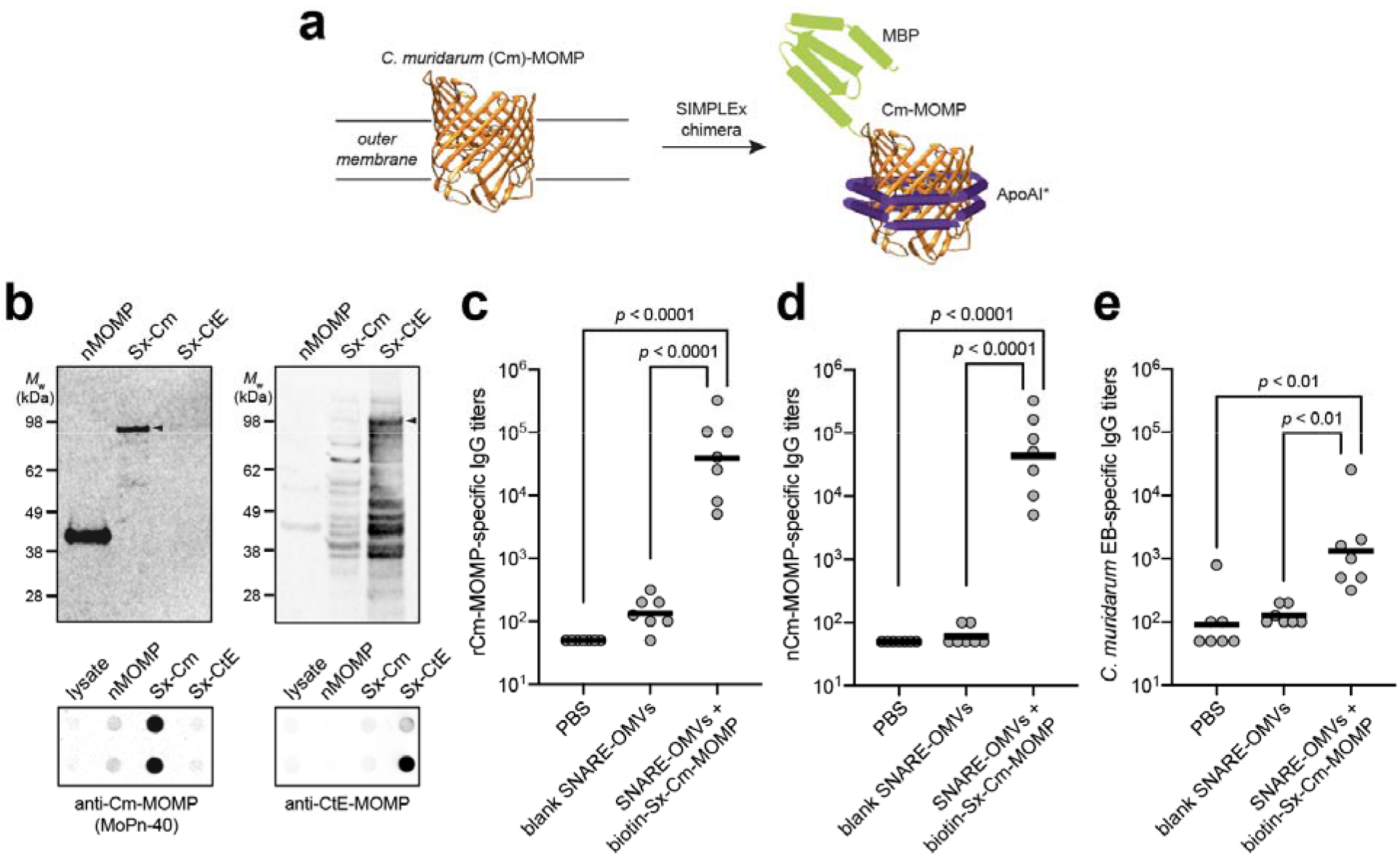
SNARE-OMVs decorated with biotinylated Sx-Cm-MOMP elicit pathogen-specific IgGs. (a) Schematic of SIMPLEx strategy for converting integral membrane proteins into water-soluble proteins that can be expressed at high titers in the cytoplasm of host cells. Here, the β-barrel outer membrane protein Cm-MOMP was fused at its N-terminus with *E. coli* maltose-binding protein (MBP) and at its C-terminus with truncated ApoAI (ApoAI*). Structural analysis indicates that ApoAI* adopts a belt-like conformation around the membrane helices of proteins to which it is fused, effectively shielding these highly hydrophobic segments from water (*54*). (b) (left blot) Antigenicity of Sx-Cm-MOMP construct evaluated by immunoblot analysis using mAb MoPn-40. Native Cm-MOMP (nCm-MOMP) served as a positive control while Sx-CtE-MOMP served as a negative control. (right blot) The latter construct was detected with commercial antibody specific for CtE-MOMP, which did not react with Sx-Cm-MOMP or nMOMP. Expected location of full-length SIMPLEx fusion proteins are denoted by black arrows. Molecular weight (*M*_w_) ladder is indicated at left. (c) Total IgG titers against recombinant preparations of Cm-MOMP (rCm-MOMP) in endpoint (day 56) serum of individual mice (gray dots) and median titers of each group (horizontal black lines). Three groups of BALB/c mice, seven mice per group, immunized s.c. with the following: PBS, SNARE-OMVs isolated from KPM404 Δ*nlpI* cells expressing the Lpp-OmpA-eMA construct, and SNARE-OMVs mixed with biotinylated Sx-Cm-MOMP. Mice received prime injections containing an equivalent amount of OMVs (20 μg total protein) on day 0 and were boosted on day 21 and 42 with the same doses. (d, e) Same as in (c) but with either (d) a native preparation of Cm-MOMP (nCm-MOMP) or (e) elementary bodies (EBs) as immobilized antigens. Statistical significance of antibody titers for SNARE-OMVs + biotin-Sx-Cm-MOMP against blank SNARE-OMVs and PBS indicates statistically significant differences (*p* < 0.0001 for ELISAs with rCm-MOMP and nCm-MOMP; *p* < 0.01 for ELISA with EBs; unpaired *t* test with Welch’s correction) between the groups.

Cm-MOMP (*67*). We observed that mAb MoPn-40 specifically recognized the water-soluble Sx-Cm-MOMP construct but not a SIMPLEx control construct comprised of different MOMP from *C. trachomatis* serovar E (Sx-CtE-MOMP) in both denature immunoblots and non-denatured dot blots (**Fig. 5b**), indicating that water-soluble Sx-Cm-MOMP retained conformational antigenicity. Next, BALB/c mice were immunize s.c. with SNARE-OMVs decorated with biotinylated Sx-Cm-MOMP. When the resultin immune sera was analyzed for reactivity against either a recombinant or nativ preparation of Cm-MOMP (rCm-MOMP and nCm-MOMP, respectively) (*51*), w observed strong cross-reaction to both antigens with total IgG titers that wer significantly greater (*p* < 0.0001) than the titers elicited by blank SNARE-OMVs an PBS control groups (**Fig. 5c and d**). It is also worth noting that the IgG responses triggered by Sx-Cm-MOMP docked on SNARE-OMVs were *Chlamydia-*specific as evidenced by the binding to *C. muridarum* elementary bodies (EBs), which was significantly above the binding measured for blank SNARE-OMVs and PBS control groups (**Fig. 5e**). As expected, antibody titers to rCm-MOMP and nCm-MOMP were similar while titers to EBs were lower. It should be pointed out that comparing titers between MOMP and EBs is not possible because the amount of MOMP present in EBs was not quantitated. Nonetheless, these data are significant because they demonstrate that the immune system of the mouse was able to recognize Cm-MOMP in the context of an OMV-tethered SIMPLEx construct. Taken together, these results confirm that dock-and-display of SIMPLEx-solubilized variants of membrane proteins on SNARE-OMVs is a unique approach for rapidly engineering vaccines based on difficult-to-obtain membrane-bound protein antigens without compromising antigenicity or immunogenicity.

## Discussion

In this study, we have developed a universal platform called AddVax for rapidly assembling antigens of interest on the surface of OMVs. The method involves site-specific docking of biotinylated antigens to the exterior of ready-made OMVs displaying multiple copies of highly modular receptors called SNAREs, which are engineered by fusing an outer membrane scaffold domain to a biotin-binding domain. As we showed here, SNARE-OMVs can be readily adorned with virtually any antigen that is amenable to biotinylation including globular and membrane proteins, glycans and glycoconjugates, haptens, lipids, and short peptides. The ability to precisely and homogenously load OMVs with a molecularly diverse array of subunit antigens differentiates the AddVax method from previous covalent conjugation strategies that are largely restricted to protein and peptide antigens (*31, 33, 34*). Moreover, our dock-and-display approach side-steps many of the challenges associated with display on OMVs using conventional genetic fusion and cellular expression technology, thereby opening the door to important vaccine subunit antigens such as malarial Pfs25 and *Chlamydia* Cm-MOMP that are refractory to soluble expression and outer membrane localization in *E. coli* (*45, 50-53*). While the separate preparation of a biotinylated antigen adds an extra step, it affords an opportunity to generate protein antigens using different expression systems, which can be chosen based on their ability to promote high yields and desired conformations including post-translational modifications.

When injected in wild-type BALB/c mice, SNARE-OMV formulations displaying GFP or a water-soluble variant of Cm-MOMP were capable of triggering strong antigen-specific humoral responses that depended on the physical linkage between the antigen and the SNARE-OMV delivery vehicle. Importantly, the GFP-specific IgG titers elicited by GFP-studded SNARE-OMVs rivaled that of ClyA-GFP-containing OMVs generated by conventional cellular expression technology (*22*). This ability of SNARE-OMVs to amplify the immunogenicity of GFP, a weakly immunogenic protein by itself (*22, 65*), without the need for potentially reactogenic adjuvants indicates that the inbuilt adjuvanticity of OMVs is preserved in the context of our dock-and-display strategy. In the case of the validated subunit vaccine candidate, Cm-MOMP (*49, 51*), we demonstrate the potential of AddVax to be readily combined with SIMPLEx, a technology for solubilizing integral membrane proteins (*54, 68*), leading to an entirely new strategy for formulating difficult-to-obtain antigens without compromising immunogenicity. Future adaptations of AddVax could also be pursued as needed such as increasing antigen density with tandemly repeated biotin-binding modules or enabling multi-antigen display with SNAREs comprised of multiple orthogonal protein-ligand binding pairs. Along these lines, we previously engineered a trivalent protein scaffold containing three divergent cohesin domains for the position-specific docking of a three-enzyme cascade on the exterior of OMVs (*69*), which provides a conceptual starting point for next-generation SNARE-OMVs.

The AddVax technology is based on the extraordinarily high affinity of avidin for the small molecule biotin and was found to be compatible with a range of different biotin-binding modules including eMA, RA, and mSA^S25H^. Although the binding affinity of our preferred biotin-binding domain, eMA, toward free biotin is measurably weaker than tetrameric SA (*K*_d_ = 31 x 10^−12^ M for eMA versus ~10^−14^ M for SA), eMA is reported to have almost multimeric avidin-like binding stability toward biotin conjugates (*38*), making it an incredibly useful module for capturing diverse subunit antigens as we showed here. Moreover, its small, monomeric design resulted in significantly better expression and OMV localization of the Lpp-OmpA-eMA SNARE compared to Lpp-OmpA-SA, which in turn resulted in far superior antigen capture. Another notable trait of eMA is its ability to be stored at −20°C without visible aggregation or loss of binding function (*38*), which could prove useful in the future for long-term vaccine storage. It should be noted that the versatility of the avidin-biotin technology has been previously leveraged as a building material in other types of vaccine formulations, enabling the attachment of antigens onto virus-like particles (VLPs) (*70–72*) and the self-assembly of macromolecular complexes comprised of vaccine antigens (*73, 74*). However, to our knowledge, our study is the first to repurpose avidin-biotin for antigen self-assembly and display on OMVs.

Overall, the AddVax platform enables creation of antigen-studded OMVs with the potential to impact many important facets of vaccine development. For example, the simplicity and modularity of vaccine self-assembly using AddVax enables rapid cycles of development and testing, which could be useful for evaluating large numbers and different combinations of pathogen-derived antigens for their ability to combat the most intractable diseases such as malaria or tuberculosis. Moreover, the fact that AddVax is based on an identical, easy-to-decorate SNARE-OMV scaffold that can be readily mass produced could shorten the time from development to manufacturing and accelerate regulatory review for each new vaccine candidate. The universal scaffold also affords the ability to share production costs across multiple antigens and diseases, which in combination with the favorable manufacturing economics of *E. coli-*based production, could help to meet the target of US $0.15 per human vaccine dose set by the Bill and Melinda Gates Foundation. In addition, pre-production of modular OMV scaffolds that can be stably stored at −20°C and then only need to be mixed with good manufacturing practice (GMP)-grade biotinylated antigens could enable rapid responses to pathogen outbreaks or pandemics. One major remaining obstacle is the fact that OMVs derived from laboratory strains of *E. coli* have yet to enter the clinic. It should be noted, however, that OMVs/OMPCs from *Neisseria meningitidis* serogroup B are the basis of two licensed vaccines, PedvaxHIB^®^ and Bexsero^®^, that are approved for use in humans (*15, 16*). Hence, although more testing of SNARE-OMV vaccine candidates is clearly required, including broader immunogenicity testing and pathogen challenge studies, we anticipate that clinical translation may not be far off.

## Materials and Methods

### Strains, growth media, and plasmids

All OMVs in this study were isolated from the hypervesiculating *E. coli* strain KPM404 Δ*nlpI* (*40*), which contains several genetic modifications that render its LPS less reactogenic. *E. coli* strain BL21(DE3) (Novagen) was used to express GFP and rCm-MOMP. The SIMPLEx constructs Sx-Cm-MOMP and Sx-CtE-MOMP were produced in two different ways, cell-based expression with BL21(DE3) or cell-free expression using an *E. coli*-based translation kit (RTS 500 ProteoMaster *E. coli* HY kit, Biotechrabbit GmbH) as described previously (*75*). Both methods yield comparable amounts of similar quality products as assessed by SDS-PAGE and immunoblot analysis. *E. coli* strain CLM24 was used to produce CRM197 conjugated with *Ft*O-PS (*58*) while strain JC8031 (*76*) was used to produce recombinant *Ft*LPS. Recombinant GFP used for serum antibody titering was expressed and purified from *Saccharomyces cerevisiae* strain SEY6210.1 to avoid cross-reaction of serum antibodies with contaminating host proteins present in protein preparations derived from *E. coli* cultures. The *C. muridarum* strain Nigg II (ATCC VR-123) was used to produce nCm-MOMP and EBs utilized in serum antibody titering experiments.

For cloning and strain propagation, *E. coli* strains were grown on solid Luria-Bertani LB (10 g/L tryptone, 10 g/L NaCl, and 5 g/L yeast extract) supplemented with agar (LBA) and yeast strain SEY6210.1 was grown on synthetic defined media without uracil (SD-URA; MP Biomedicals) supplemented with agar. For OMV production, hypervesiculating *E. coli* were grown in terrific broth (TB) (12 g/L tryptone, 24 g/L yeast extract, 0.4% v/v glycerol, 0.17 M KH_2_PO_4_ and 0.72 M K_2_HPO_4_). For production of recombinant antigens using *E. coli*, cells were grown in LB media. SEY6210.1 was grown in SD-URA or yeast extract-peptone-dextrose (YPD) media (20 g/L peptone, 10 g/L yeast extract and 2% w/v glucose). *C. muridarum* cells were grown as described (*77*) (*78*).

All plasmids used in this study are described in **Supplementary Table 1**. Standard restriction enzyme-based cloning methods were used and sequences were confirmed through Sanger sequencing performed by the Cornell Biotechnology Resource Center (BRC) unless specified otherwise. For expression of SNARE constructs in OMVs, eMA fusions to ClyA, Lpp-OmpA, and the membrane-associated transporter domains of the autotransporters Int, Hbp, Ag43, and IgAP were codon-optimized for *E. coli* expression, synthesized, and cloned into plasmid pBAD24 (*79*) between EcoRI and SphI restriction sites with an NdeI site at the start codon by GenScript. SNARES involving ClyA, Lpp-OmpA and Int were cloned with eMA fused to the 3’ end of the scaffold while SNAREs involving Hbp, Ag43 and IgAP were cloned with eMA fused to the 5’ end of the scaffold (**Fig. 1b**). For the latter set of constructs, DNA encoding a Sec-dependent export signal peptide derived from PelB (spPelB), identical to the sequence in pET22b (Novagen), was introduced at the 5’-end of the eMA-scaffold gene fusions. For all of these constructs, DNA encoding c-Myc (EQKLISEEDL) and FLAG (DYKDDDDK) epitope tags was introduced at the 5’ and 3’ ends of eMA as depicted in **Fig. 1b**. In the case of the autotransporter AIDA-I, the transporter unit (amino acids 962 through 1286) was PCR-amplified from pIB264 (*80*) and ligated into pBAD24 between XhoI and SphI restriction sites. A “gBlock” (Integrated DNA Technologies, IDT) encoding eMA with a 5’ FLAG tag (IDT) was ligated between NcoI and XhoI restriction sites, after which a gBlock encoding spPelB (IDT) was ligated between EcoRI and NcoI.

To construct L-rhamnose inducible plasmids, DNA encoding Lpp-OmpA-eMA and eMA-IgAP was digested from the respective pBAD24 expression vectors and ligated into pTrham (Amid Biosciences) between NdeI and SphI sites, yielding plasmids pTrham-Lpp-OmpA-eMA and pTrham-eMA-IgAP, respectively. To construct SNAREs based on alternative avidin domains, *Rhizobium etli* RA, *Streptomyces avidinii* SA, and an optimized version of monomeric streptavidin, namely mSA^S25H^, with a lowered off-rate (*43*) were codon-optimized and synthesized as gBlocks with flanking BbsI and HindIII restriction sites (IDT). The sequences were then used to replace the eMA sequence in the pTrham-Lpp-OmpA-eMA vector, resulting in plasmids pTrham-Lpp-OmpA-RA, pTrham-Lpp-OmpA-SA, and pTrham-Lpp-OmpA-mSA.

For expression of GFP antigen for docking on OMVs, the gene encoding FACS-optimized GFPmut2 with a C-terminal 6xHis tag was cloned in pET24a(+)-Cm^R^ between SacI and HindIII restriction sites, yielding pET24-GFP. For yeast expression of GFP used in serum antibody titering, a codon-optimized gene encoding GFPmut2 was synthesized as a double-stranded DNA fragment or gBlock (IDT) with a 5’ Kozak sequence and 3’ 6xHis tag and ligated into the yeast-expression plasmid pCM189 (ATCC) between BamHI and PstI sites, yielding pCM-GFP. For expression of the Sx-Cm-MOMP antigen for OMV docking studies, the sequence encoding codon-optimized Cm-MOMP, which was designed previously (*75*), was synthesized as a gBlock (IDT) and ligated into the SIMPLEx plasmid pET21d-Sx (*68*) between NdeI and EcoRI restriction sites, yielding pET21-Sx-Cm-MOMP. A modified strategy was used to generate plasmid pIVEX-Sx-CtE-MOMP encoding the Sx-CtE-MOMP construct. Briefly, codon optimized CtE-MOMP was generated in-house following a previously described strategy for Cm-MOMP (*75*). PCR products corresponding to CtE-MOMP, MBP, and ApoAI* (human ApoAI with 49 N-terminal amino acids removed) were cloned into pIVEX-2.4d using the following restriction enzyme strategy: NdeI-MBP-XhoI-MOMP-NsiI-ApoA1-SacI. The plasmid sequence was confirmed through Sanger sequencing performed by ElimBiopharm.

### Protein purification

For production of GFP and Sx-Cm-MOMP protein antigens, BL21(DE3) cells containing plasmids corresponding to each antigen were grown overnight at 37°C in 5 mL LB supplemented with the appropriate antibiotic and subcultured 1:100 into the same media. Protein expression was induced with 0.1 mM isopropyl-β-D-1-thiogalactopyranoside (IPTG) when culture densities reached an absorbance at 600 nm (Abs_600_) of ~1.0 and proceeded for 16 h at 30°C. Cells were then harvested and lysed by homogenization, and proteins were purified by Ni-NTA resin (Thermo-Fisher) following the manufacturer’s protocol. For Sx-Cm-MOMP, Ni-NTA resin elute was immediately diluted with PBS containing 1 mM EDTA (PBS-E) and incubated with amylose resin (New England Biolabs) for 30 min, followed by washing with 10 resin volumes of PBS-E and elution with 10 mM maltose in PBS-E. All purified proteins were buffer exchanged into PBS using PD-10 desalting columns (Cytiva), filter-sterilized, quantified by Lowry (MilliporeSigma), and stored at 4°C for up to 2 months or at −80°C for longer term storage. Pfs25 was expressed and purified from a baculovirus expression system using *Spodoptera frugiperda* SF9 cells and P2 virus by Genscript.

To produce CRM197-*Ft*O-PS glycoconjugate, *E. coli* strain CLM24 was transformed with plasmid pTrc99S-spDsbA-CRM197^4xDQNAT^ encoding the CRM197 carrier protein modified at its C-terminus with four tandemly repeated DQNAT glycosylation motifs (*81*), plasmid pGAB2 encoding the *Ft*O-PS biosynthesis pathway (*58*), and plasmid pMAF10-PglB encoding the *Campylobacter jejuni* oligosaccharyltransferase PglB for transfer of the *Ft*O-PS (*82*). Overnight cultures were subcultured 1:100 into fresh LB containing appropriate antibiotics. When culture densities reached Abs_600_ of ~0.8, PglB expression was induced with 0.2% arabinose for 16 h at 30°C, at which point CRM197^4xDQNAT^ expression was induced with 0.1 mM IPTG and cells were grown for an additional 8 h at 30°C. Cells were then harvested and purified as described above for GFP.

To purify GFP for serum antibody titering, yeast strain SEY6210.1 was transformed with pCM189-GFP-6xHis and grown on SD-URA agar plates at 30°C for two days. Afterwards, a colony was picked and grown overnight at 30°C in 5 mL of SD-URA media containing tetracycline, subcultured 1:10 into YPD, and grown for 20 h at 30°C. Yeast cells were lysed by homogenization and protein was purified by Ni-NTA resin as above. For Cm-MOMP serum antibody titering, rCm-MOMP was expressed recombinantly in *E. coli* while nCm-MOMP was extracted from *C. muridarum* strain Nigg II as described previously (*51*).

### Antigen biotinylation

Purified GFP, Pfs25, Sx-Cm-MOMP, and CRM197-*Ft*O-PS were mixed at 1 mg/mL (0.25-1 mg total protein) with 1.5x molar excess EZ-Link Sulfo-NHS-LC biotin (Thermo-Fisher) in PBS and incubated on ice for 2-3 h. Afterwards, the reaction mix was passed five times over PBS-equilibrated monomeric avidin resin (Thermo-Fisher). Final flow-through fractions were concentrated and saved for repeat biotinylation reactions as needed. Following 6 washes each with one resin volume of PBS, biotinylated protein was eluted 6 times each with one resin volume of 2 mM D-biotin (MilliporeSigma) in PBS. Elutions were pooled and diluted to 6 mL with PBS and concentrated to <200 μL using 6-mL, 10-kDa cut-off protein concentrators (Pierce). Dilution and concentration was repeated three more times to remove the D-biotin. The final concentrated biotinylated proteins were filter-sterilized, quantified by Lowry, and stored at 4°C for up to 2 months.

To biotinylate *F. tularensis* LPS, we first purified *Ft*LPS as described previously (*28*) with the addition of DNase-I (0.5 mg/mL; MilliporeSigma) in the Proteinase K treatment step. To remove sugar monomers and short polysaccharide chains, *Ft*LPS was buffer exchanged into PBS using PD-10 columns and quantified by the Purpald assay (*83*). Biotinylation was performed as described previously using 1-cyano-4-dimethylaminopyridinium tetrafluoroborate (CDAP) as the activation reagent linking EZ-Link-Amine-PEG3-Biotin (Pierce) to hydroxyl groups on the polysaccharide (*73*). A 27-amino acid B16-M30 peptide with N-terminal biotin and C-terminal polyhistidine (6xHis) motif for antibody-based detection was synthesized by Biomatik to ~85% purity, and a 1 mg/mL stock was prepared in dimethyl sulfoxide (DMSO). Biotinylated GD2 ganglioside oligosaccharide and biotinylated Le^Y^ oligosaccharide were purchased from Elicityl, biotinylated DNP containing a polyethylene glycol (PEG) linker was purchased from Nanocs, and 1-oleoyl-2-[12-biotinyl(aminododecanoyl)]-sn-glycero-3-phosphocholine (18:1-12:0 biotin PC) powder was purchased from Avanti Polar Lipids. The biotin-GD2, biotin-Le^Y^, and biotin-DNP were dissolved in sterile water (1-5 mg/mL) while biotin-PC was suspended in DMSO (1 mg/mL). All stocks were diluted in PBS for avidin binding studies.

### OMV preparation

KPM404 Δ*nlpI* cells containing pBAD24 or pTrham expression plasmids were spread from −80°C glycerol stocks onto LBA plates supplemented with 100 μg/ml carbenicillin and grown overnight at 37°C (~20 h). On the following day, cells were suspended from the agar using TB and subcultured to Abs_600_ of ~0.06 in 50-100 mL TB supplemented with carbenicillin. Cells were grown at 37°C and 220 rpm and induced when Abs_600_ reached ~0.6 to ~1.8 with varying concentrations of L-arabinose (pBAD24) or L-rhamnose (pTrham). Following induction, cells were grown for 16 h at 28°C followed by 6 h at 37°C, after which cells were pelleted via centrifugation at 10,000 x g for 15 min. Supernatants were filtered through 0.2 μm filters and stored overnight at 4°C. OMVs were isolated by ultracentrifugation at 141,000 x g for 3 h at 4°C and resuspended in sterile PBS. For quantitative analysis and immunizations, resuspended OMVs were diluted in sterile PBS and ultracentrifuged a second time to remove residual media and soluble proteins. Following a second resuspension in PBS, large irreversible aggregates were removed by centrifuging for 2 min at 3,000 x g in a microcentrifuge and filtering the supernatant using sterile 4-mm, 0.45-μm syringe filters (MilliporeSigma). Total OMV proteins were quantified by Lowry (Peterson’s modification; MilliporeSigma) using bovine serum albumin (BSA) as protein standard. OMVs were stored for up to 1 month at 4°C for binding analysis and up to 2 weeks prior to immunizations.

### Immunoblot analysis

Biotinylated and non-biotinylated protein antigens and OMVs were mixed with loading buffer containing β-mercaptoethanol and boiled for 10 min prior to loading onto Mini-PROTEAN TGX polyacrylamide gels (Bio-Rad). To determine protein purity, gels were stained with Coomassie G-250 stain (Bio-Rad) following the manufacturer’s protocol. For immunoblot analysis, proteins were transferred to polyvinylidene difluoride (PVDF) membranes and blocked with 5% milk followed by probing with antibodies, which were all used at 1:5,000 dilution. Avidin expression on OMVs was analyzed with horseradish peroxidase (HRP)-conjugated anti-c-Myc (Abcam; Cat # ab19312) or HRP-conjugated anti-DDDDK (Abcam; Cat # ab1162) antibodies that recognized c-Myc and FLAG epitope tags, respectively. Proteins and peptides bearing C-terminal 6xHis tags were detected with mouse anti-6xHis antibody clone AD1.1.10 (BioRad; Cat # MCA1396GA) while detection of glycosylated CRM197-*Ft*O-PS was with anti-*F. tularensis* LPS antibody clone FB11 (Invitrogen; Cat # MA1-21690) that is specific to *Ft*LPS (*28*). HRP-conjugated goat anti-mouse (Abcam; Cat # ab6789) was used as needed. All membranes were developed using Clarity ECL substrate (Bio-Rad) and visualized using a ChemiDoc imaging system (Bio-Rad).

For probing antigenicity of SIMPLEx constructs, Sx-Cm-MOMP and Sx-CtE-MOMP were mixed with loading buffer containing DTT and boiled for 10 min before loading onto 4-12% NuPAGE Bis-Tris gels (Thermo-Fisher). For denaturing immunoblot analysis, proteins were transferred to PVDF membranes and blocked with 5% BSA (Sigma) followed by probing with mAb MoPn-40 (1:1,000) (*67*) or anti-CtE-MOMP (1:2,000; Novus Biologicals; Cat # NB100-66403) antibodies. For non-denaturing dot blot analysis, purified Sx-Cm-MOMP and Sx-CtE-MOMP proteins were spotted directly onto nitrocellulose membranes and incubated for 5 min before being blocked with 5% BSA (Sigma) and probing with the same primary antibodies. IRDye 800CW-conjugated goat anti-mouse secondary antibodies (1:10,000; Li-Cor; Cat # 926-32210) were used to detect primary antibody binding and membranes were visualized using an Odyssey Li-Cor Fc imaging system.

### TEM analysis of OMVs

Ultrastructural analysis of OMVs was performed via TEM as previously described (*22*) with a few modifications. Briefly, OMVs were diluted to 100 μg/mL and negatively stained with 1.5% uranyl acetate and deposited on 300-mesh Formvar carbon-coated copper grids. Imaging was performed using a FEI Tecnai 12 BioTwin transmission electron microscope.

### ELISA

For qualitative assessment of antigen binding by SNARE-OMVs, OMV samples were diluted to 2 μg/mL in PBS, coated on Costar 9018 high-binding 96-well plates (50 μL per well), and incubated overnight at 4°C. Plates were blocked with 2% BSA in PBS (100 μL/well) for 3 h at room temperature and subsequently washed two times with PBST (PBS pH 7.4 with 0.005% Tween-20 and 0.3% BSA). To analyze relative binding capacities, biotinylated or unbiotinylated antigens were serially diluted in triplicate by a factor of 3 in PBST, starting from 10 nM, and incubated for 90 min at room temperature (50 μL/well). Unbound antigen was removed by washing twice with PBST. Bound antigen was labeled by incubating with primary antibody for 1 h in PBST followed by two more PBST washes and a 1-h incubation with HRP-conjugated secondary antibody. After three final washes with PBST, 3,3’-5,5’-tetramethylbenzidine substrate (1-Step Ultra TMB-ELISA; Thermo-Fisher) was added and the plate was incubated at room temperature for 30 min in the dark. The reaction was stopped with 2M H_2_SO_4_ and absorbance was measured via microplate spectrophotometer (Molecular Devices) at Abs_450_. The absorbance reading from OMVs incubated with PBST without antigen was subtracted from the signal in all wells with antigen added. The resulting values were normalized to the highest average absorbance value among all antigen concentrations, including unbiotinylated antigen controls. Primary anti-6xHis and anti-*Ft*LPS antibodies and HRP-conjugated anti-mouse secondary antibody were identical to those used for immunoblotting above and were used at the same dilutions. The remaining antigens were detected with the following antibodies: GD2 was detected with mouse anti-ganglioside GD2 antibody (1:1,000; Abcam; Cat # ab68456); Le^Y^ was detected with mouse anti-Lewis Y antibody clone H18A (1:1,000; Absolute Antibody; Cat # Ab00493-1.1); DNP was detected with goat anti-DNP (1:5,000; Bethyl Laboratories; Cat # A150-117A); and PC was detected with anti-phosphorylcholine antibody clone BH8 (1:250; MilliporeSigma; Cat # MABF2084). HRP-conjugated donkey anti-goat secondary was used as needed (1:5,000; Abcam; Cat # ab97110).

For quantification of antigen binding capacity on SNARE-OMVs, 50 μg OMVs were diluted to 0.1 mg/mL in PBS, mixed with unbiotinylated or biotinylated GFP at concentrations between 0 and 100 pmol/mL (0 to 1 pmol antigen/μg OMV), and incubated at room temperature for 1 h. Mixtures were then diluted to 30 mL in PBS and ultracentrifuged for 141,000 x g for 3 h at 4°C. After discarding the supernatant, the pellet was resuspended with 100 μL PBS, and the washed OMVs were quantified by Lowry (Peterson’s modification). An ELISA-based method that could be applied to a variety of molecules was then used to quantify the amount of antigen remaining. Specifically, washed OMVs were diluted to 2 μg/mL and coated on high-binding ELISA plates (Costar 9018) in triplicate (50 μL per well). Known standards were prepared by mixing blank SNARE-OMVs at 2 μg/mL with 1:2 serial dilutions of unbiotinylated or biotinylated GFP, starting from 1 pmol GFP/μg OMV, and coating each antigen concentration in triplicate on the ELISA plates (50 μL per well). Following overnight coating at 4°C, plates were blocked with 2% BSA in PBS for 3 h (100 μL per well) at room temperature and subsequently washed two times with PBST. Antibody and substrate incubations were identical to the qualitative binding ELISA described above. The final Abs_450_ signals of the OMV mixtures containing known amounts of unbiotinylated or biotinylated antigen were used to generate standard curves from which the amount of antigen remaining in each unknown washed OMV sample was calculated. The amount of GFP displayed on positive-control ClyA-GFP OMVs was quantified by fluorescence as described previously (*22*).

### Mouse immunizations

One day prior to immunization (day −1), different OMV formulations were diluted to 0.1 mg/mL in sterile PBS pH 7.4. For the formulations involving docked antigens, antigens and OMVs were mixed to a final concentration of 100 pmol/mL and 0.1 mg/mL, respectively, corresponding to 1 pmol antigen/μg OMV (~3 wt% for GFP). All formulations were immediately stored at 4°C. On day 0, 200 μL (20 μg) OMVs were injected subcutaneously into six-week-old BALB/c mice (7 mice per group). Identical booster injections were administered on days 21 and 42, and blood was drawn from the mandibular sinus on days −1, 28 and 49. Mice were euthanized on day 56, which was immediately followed by blood collection via cardiac puncture and spleen collection. The protocol number for the animal trial was 2012-0132 and was approved by the Institutional Animal Care and Use Committee (IACUC) at Cornell University.

### Serum antibody titers

Sera was isolated from the blood of immunized mice by centrifugation at 5,000 x g for 10 min and stored at −20°C. Antigen-specific antibodies in the sera were measured using indirect ELISA as described previously (*28*) with a few modifications. Briefly, high-binding 96-well plates (Costar 9018) were coated with purified antigen (5 μg/mL in PBS, pH 7.4) and incubated overnight at 4°C, followed by overnight blocking with 5% non-fat dry milk (Carnation) in PBS. Serum samples were serially diluted in triplicate by a factor of two in blocking buffer, starting from 1:100, and incubated on the antigen-coated plates for 2 h at 37°C. Plates were washed 3 times with PBST and incubated for 1 h at 37°C in the presence of one of the following HRP-conjugated antibodies: goat anti-mouse IgG (1:10,000; Abcam Cat # ab6789); anti-mouse IgG1 (1:10,000; Abcam Cat # ab97240), or anti-mouse IgG2a (1:10,000; Abcam Cat # ab97245). Following 3 final washes with PBST, 1-Step Ultra TMB-ELISA (Thermo-Fisher) was added and the plate was incubated at room temperature for 30 min in the dark. The reaction was stopped with 2M H_2_SO_4_, and absorbance was quantified via microplate spectrophotometer (Molecular Devices) at Abs_450_. Serum antibody titers were determined by measuring the highest dilution that resulted in signal three standard deviations above no-serum background controls.

The *Chlamydia*-specific antibody titers in sera from mice immunized with Sx-Cm-MOMP were determined by ELISA as previously described (*51*). Briefly, 96-well plates were coated with 2 μg/ml of rCm-MOMP or nCm-MOMP, or 100 μL/well of *C. muridarum* EBs containing 10 μg/mL of protein in PBS. Next, 100 μL of serum was added per well in serial dilutions. Following incubation at 37°C for 1 h, the plates were washed, and HRP-conjugated goat anti-mouse IgG (1:10,000; BD Biosciences Cat # 554002) was added. The plates were incubated and washed, and the binding was measured in an ELISA reader (Labsystem Multiscan) using 2,2′-azino-bis-(3-ethylbenzthiazoline-6-sulfonate) as the substrate.

### Statistical analysis

Statistical significance between groups was determined by unpaired *t*-test with Welch’s correction using GraphPad Prism software (version 9.0.2). Statistical parameters including the definitions and values of *n*, *p* values, and SDs are reported in the figures and corresponding figure legends.

## Supporting information

Supplementary Results, Table and Figures

## Data availability.

All data needed to evaluate the conclusions in the paper are present in the paper and/or the Supplementary Information.

## Acknowledgements.

We thank the PATH Malaria Vaccine Initiative (MVI) for providing the Pfs25 construct. This work was supported by the Bill and Melinda Gates Foundation (OPP1217652), the Defense Threat Reduction Agency (grant HDTRA1-20-10004), the National Institutes of Health (grants R01GM137314 and R01GM127578 to M.P.D. and U19AI144184 to S.P., L.M.M., and M.A.C.), and the National Science Foundation (grants CBET-1605242 and CBET-1936823 to M.P.D.). The work was also supported by seed project funding (to M.P.D.) through the National Institutes of Health-funded Cornell Center on the Physics of Cancer Metabolism (supporting grant # U54CA210184-01). The content is solely the responsibility of the authors and does not necessarily represent the official views of the National Cancer Institute or the National Institutes of Health. K.B.W. was supported by a National Science Foundation Graduate Research Fellowship (grant DGE-1650441) and a Cornell Fleming Graduate Scholarship. T.J. was supported by a Royal Thai Government Fellowship and a Cornell Fleming Graduate Scholarship. T.D.M. was supported by a training grant from the National Institutes of Health NIBIB (T32EB023860) and a Cornell Fleming Graduate Scholarship.

## Author Contributions.

K.B.W. designed research, performed research, analyzed data, and wrote the paper. T.J., T.D.M., S.H.-P., S.P., S.F.G., M.R.-D.J., R.S., and J.L. designed research, performed research, and analyzed data. C.L. and D.P. wrote the paper. L.M.M., M.A.C. and M.P.D. designed and directed research, analyzed data, and wrote the paper.

