## Supplementary Results, Table and Figures for "A modular platform for on-demand vaccine self-assembly enabled by decoration of bacterial outer membrane vesicles with biotinylated antigens"

The two most effective SNAREs in terms of biotin-GFP binding, namely eMA-IgAPβ and Lpp-OmpA-eMA, were evaluated over a range of conditions to identify parameters that affected GFP docking levels. A preliminary test of different cultivation variables (e.g., growth temperature, culture density at time of induction, inducer level, plasmid backbone, etc.), revealed that the density of the culture at the time of receptor induction had the greatest impact on the levels of biotin-GFP loading, with higher induction densities ($Abs_{600}$ $\approx$ 1.8) resulting in SNARE-OMVs that captured the most antigen (**Supplementary Fig. 2a**). The Lpp-OmpA-eMA construct showed the highest biotin-GFP binding levels under the conditions tested. However, expression of this SNARE was detrimental to the host cells based on the observation that the final culture densities hardly changed, and in some cases even decreased, from the densities at the time of induction, which was not the case for IgAP-eMA (**Supplementary Fig. 2b**). Given the different biogenesis pathways of the IgAP autotransporter versus the Lpp-OmpA β-barrel outer membrane protein, we suspected that the host cell toxicity associated with Lpp-OmpA might result from inducer levels that were too strong. In support of this notion, when Lpp-OmpA-eMA constructs were induced with ~50-times less inducer (0.27 mM vs. 13.3 mM L-arabinose), the post-induction cell growth was markedly improved, with Lpp-OmpA-eMA-expressing cells reaching a final density on par with that of cells expressing IgAP-eMA (**Supplementary Fig. 2c**). Importantly, the Lpp-OmpA-eMA SNARE-OMVs isolated from these healthier host cells captured significantly more biotin-GFP compared to IgAP-eMA SNARE-OMVs.

To determine whether these effects were specific to the choice of plasmid, we evaluated an alternative plasmid for expression of both SNAREs. Specifically, we re-cloned the IgAP-eMA and Lpp-OmpA-eMA constructs into the L-rhamnose-inducible plasmid, pTrham, which is known to afford tighter expression control compared to pBAD vectors and can help to overcome the deleterious saturation of membrane and secretory protein biogenesis pathways (*1, 2*). As was observed with pBAD24, cells expressing IgAP-eMA from pTrham reached similar final densities regardless of the inducer levels, while growth of cells expressing Lpp-OmpA-eMA from pTrham decreased with increasing inducer levels (**Supplementary Fig. 2d**). Despite these differences in growth, IgAP-eMA and Lpp-OmpA-eMA SNARE-OMVs derived from cultures that were induced with 2 mM L-rhamnose each bound equivalent amounts of biotin-GFP (**Supplementary Fig. 2d**). Interestingly, increasing the amount of L-rhamnose yielded IgAP-eMA SNARE-OMVs that captured 2-3 times more biotin-GFP whereas decreasing the amount of L-rhamnose yielded Lpp-OmpA SNARE-OMVs that bound 4-5 times more biotin-GFP, consistent with the contrasting effects of inducer on the post-induction growth of cells expressing these constructs. Overall, the engineered Lpp-OmpA-eMA receptor expressed from pTrham plasmid using 0.5 mM L-rhamnose was the strongest performer in terms of biotin-GFP binding (**Supplementary Fig. 2d and e**); hence, we chose this plasmid/inducer combination for all further studies.

**Table S1.** Plasmids used in this study

| **Name** | **Description** | **Reference** |
|---|---|---|
| pET24a(+)-Cm | *E. coli* expression plasmid derived from pET-24a(+) but with $Cm^R$ resistance marker; $Cm^R$ | Lab stock |
| pET24-GFP | Encodes FACS-optimized GFPmut2 variant with C-terminal 6xHis tag in pET-24a(+)-Cm; $Cm^R$ | This study |
| pET21d-Sx | Encodes SIMPLEx components MBP at the N-terminus and ApoAI* at the C-terminus with multicloning site for insertion of POIs between MBP and ApoAI*; $Amp^R$ | (*3*) |
| pET21-Sx-Cm-MOMP | Encodes Cm-MOMP in pET21d-Sx; $Amp^R$ | This study |
| pCM189 | Yeast expression plasmid with tetracycline-regulated promoter; $Amp^R$ | (*4*) |
| pCM-GFP | Encodes *S. cerevisae* codon-optimized GFP with a C-terminal 6xHis tag in pCM189; $Amp^R$ | This study |
| pTrc99S-ssDsbA-$CRM197^{4xDQNAT}$ | Encodes DsbA signal peptide fused in-frame with CRM197 followed by a 4x tandemly repeated DQNAT glycosylation tag in pTrc99S; $Amp^R$ | (*5*) |
| pGAB2 | Encodes *F. tularensis* SchuS4 O-PS biosynthesis pathway; $Tet^R$ | (*6*) |
| pMAF10-PglB | Encodes *Campylobacter jejuni* PglB oligosaccharyltransferase in pMAF10; $Tmp^R$ | (*7*) |
| pIVEX2.4d | Plasmid for *E. coli* cell-based and cell-free expression with a strong T7 promoter; $Amp^R$ | (*8*) |
| pIVEX-Sx-CtE-MOMP | Encodes SIMPLEx fusion MBP-CtE-MOMP-ApoAI* in pIVEX2.4d; $Amp^R$ | This study |
| pET45-rCm-MOMP | Encodes Cm-MOMP without its native signal peptide in pET-45b(+); $Amp^R$ | (*9*) |
| pClyA-GFP | Encodes ClyA-GFPmut2 fusion in pBAD18-Cm; $Cm^R$ | (*10*) |
| pBAD24-ClyA-eMA | Encodes ClyA-c-Myc-eMA-FLAG fusion in pBAD24; $Amp^R$ | This study |
| pBAD24-Lpp-OmpA-eMA | Encodes Lpp-OmpA-c-Myc-eMA-FLAG fusion in pBAD24; $Amp^R$ | This study |
| pBAD24-Intimin-eMA | Encodes Int-c-Myc-eMA-FLAG fusion in pBAD24; $Amp^R$ | This study |
| pBAD24-eMA-Hbpβ | Encodes spPelB-FLAG-eMA-c-Myc-HBPβ fusion in pBAD24; $Amp^R$ | This study |
| pBAD24-eMA-Ag43β | Encodes spPelB-FLAG-eMA-c-Myc-Ag43β fusion in pBAD24; $Amp^R$ | This study |
| pBAD24-eMA-IgAPβ | Encodes spPelB-FLAG-eMA-c-Myc-IgAPβ fusion in pBAD24; $Amp^R$ | This study |
| pBAD24-eMA-AIDA-Iβ | Encodes spPelB-FLAG-eMA-AIDA-Iβ fusion in pBAD24; $Amp^R$ | This study |

| | | |
|---|---|---|
| pTrham | *E. coli* expression vector containing L-rhamnose inducible promoter rhaBAD; $Amp^R$ | Amid Biosciences |
| pTrham-Lpp-OmpA-eMA | Encodes Lpp-OmpA-c-Myc-eMA-FLAG fusion in pTrham; $Amp^R$ | This study |
| pTrham-eMA-IgAPβ | Encodes ssPelB-FLAG-eMA-c-Myc-IgAPβ fusion in pTrham; $Amp^R$ | This study |
| pTrham-Lpp-OmpA-$SA^{S25H}$ | Encodes Lpp-OmpA-c-Myc-$mSA^{S25H}$-FLAG fusion in pTrham; $Amp^R$ | This study |
| pTrham-Lpp-OmpA-SA | Encodes Lpp-OmpA-myc-SA-FLAG fusion in pTrham; $Amp^R$ | This study |
| pTrham-Lpp-OmpA-RA | Encodes Lpp-OmpA-myc-RA-FLAG fusion in pTrham; $Amp^R$ | This study |

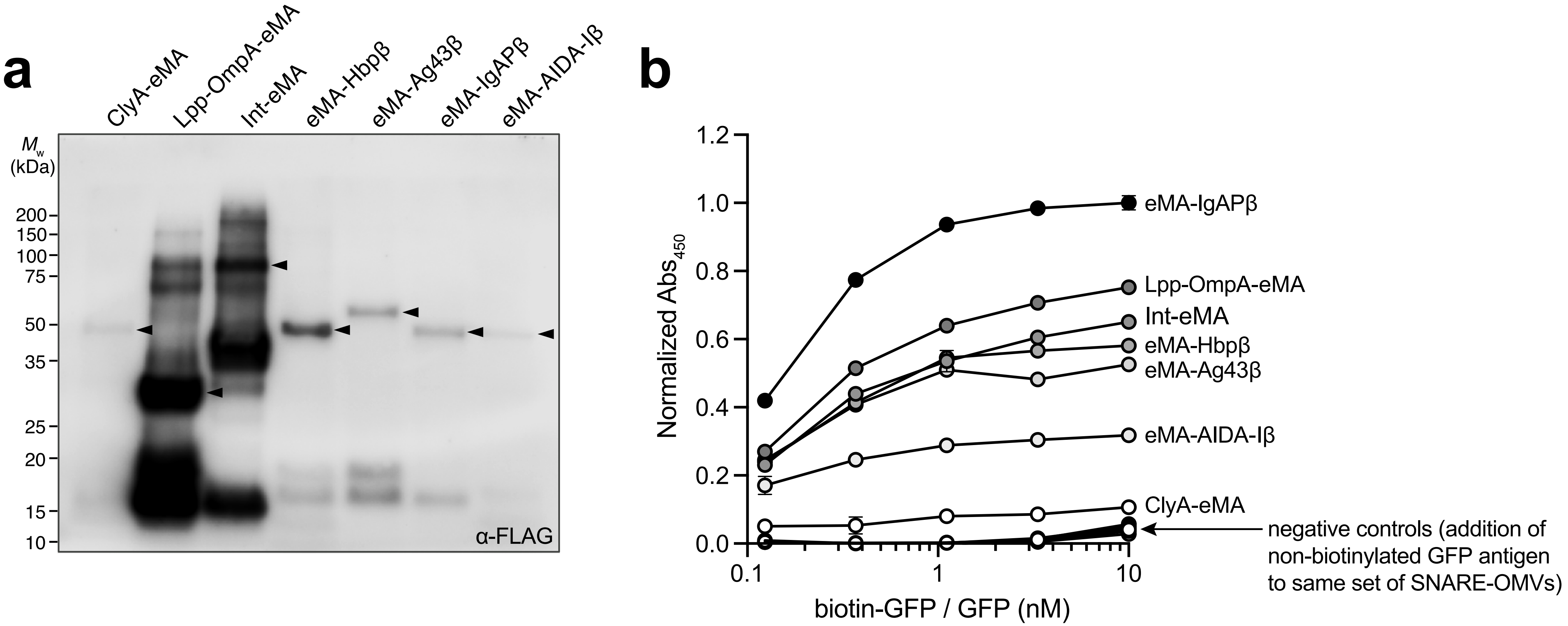


**Supplementary Figure 1. Expression and antigen-binding activity of engineered SNAREs.** (a) Immunoblot analysis of OMV fractions isolated from hypervesiculating *E. coli* strain KPM404 Δ*nlpI* expressing each of the different SNAREs from plasmid pBAD24. An equivalent amount of SNARE-OMVs as determined by total protein assay was loaded in each lane. Blot was probed with anti-FLAG antibody (α-FLAG) to detect FLAG epitope (DYKDDDDK) located at the N- or C-terminus of each construct. Expected location of full-length SNARE fusion proteins are denoted by black arrows. Molecular weight ($M_w$) ladder is indicated at left. (b) Binding of biotin-GFP to each of the different SNARE-OMVs as indicated. Binding activity was determined by ELISA in which biotin-binding SNARE-OMVs were immobilized on plates and subjected to varying amounts of biotin-GFP, after which plates were extensively washed prior to detection of bound biotin-GFP using anti-polyhistidine antibody to detect C-terminal 6xHis tag on GFP. Controls were performed by treating the same set of SNARE-OMVs with unmodfied GFP in place of biotin-GFP. All data were normalized to the maximum signal correpsonding to the eMA-IgAPβ construct in the presence of 10 nM biotin-GFP. Datapoints represent the average of three biological replicates and error bars represent the standard deviation of the mean.

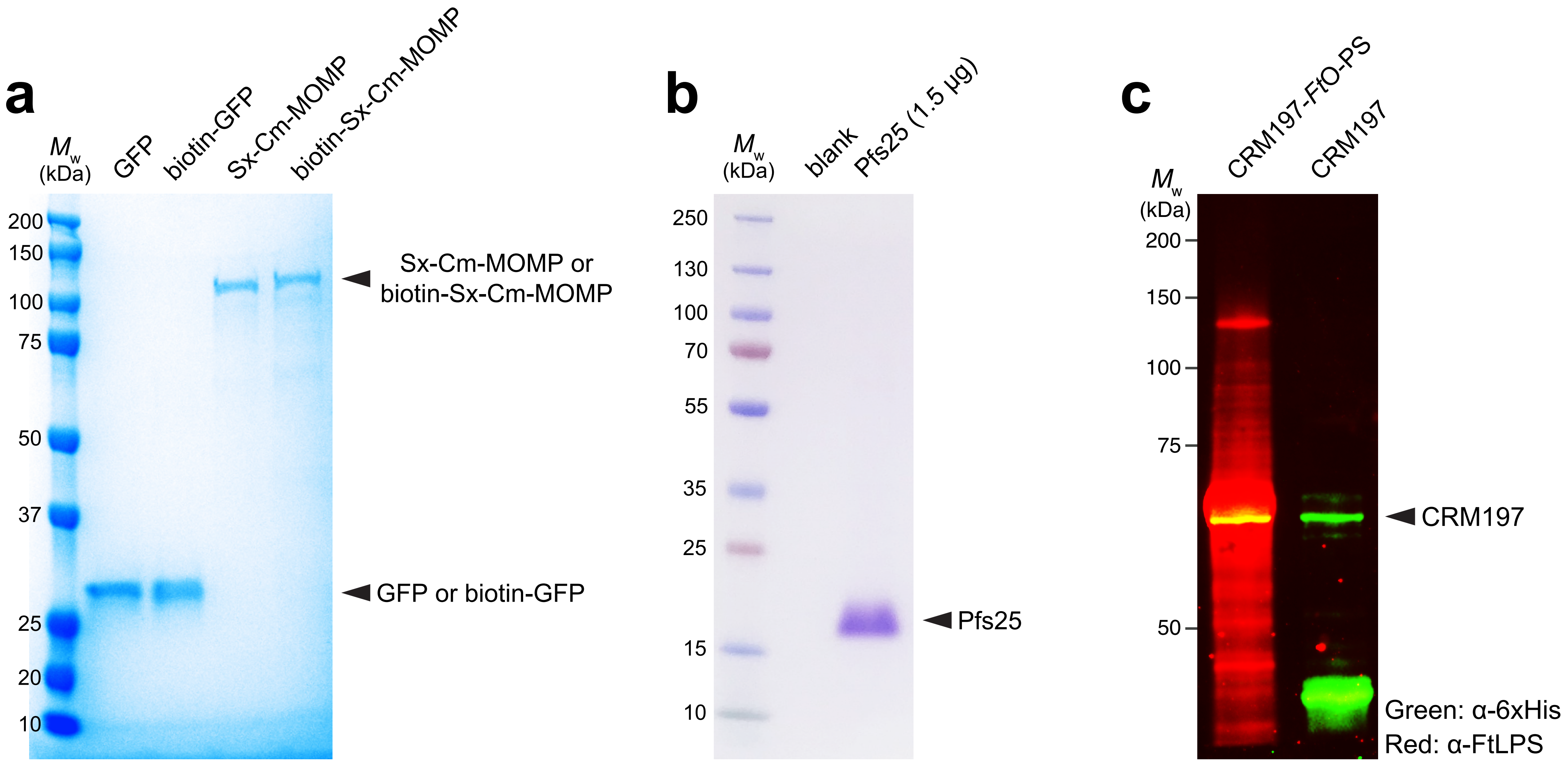


**Supplementary Figure 2. Expression and purification of biotinylated protein antigens.** Coomassie blue-stained SDS-PAGE gels of: (a) purified GFP and Sx-Cm-MOMP as well as their biotinylated counterparts; and (b) purified Pfs25. GFP and Sx-Cm-MOMP were both produced using *E. coli* BL21(DE3), which yielded ~100 mg/L of GFP and ~5 mg/L of Sx-Cm-MOMP. Note that GFP was purified by nickel only, whereas SIMPLEx-MOMP was purified by Ni-NTA resin followed by amylose. Pfs25 was produced using a baculovirus expression system involving SF9 cells and P2 virus, which yielded ~25 mg/L of Pfs25 using Ni-NTA resin. (c) Immunoblot analysis of CRM197 with four tandemly repeated DQNAT glycosylation motifs purified from *E. coli* CLM24 with (left lane) or without (right lane) plasmid DNA encoding the *Ft*O-PS biosynthetic machinery. Glyconjugate yields were typically 2-3 mg/L. Blots were probed with anti-polyhistidine antibody (α6xHis; green signal) to detect the CRM197 carrier protein or FB11 (αFtO-PS; red signal) to detect the *Ft*O-PS glycan. Image shows merge of α6x-His and α*Ft*O-PS signals. High molecular weight laddering for red signal is characteristic of variable chain length O-PS polymers that are seen in native *Ft*LPS as well as in glycoconjugates derived from engineered *E. coli* (Stark et al. 2021 *Sci Adv*). All images are representative of at least three biological replicates. Molecular weight ($M_w$) markers are shown at the left of each image.

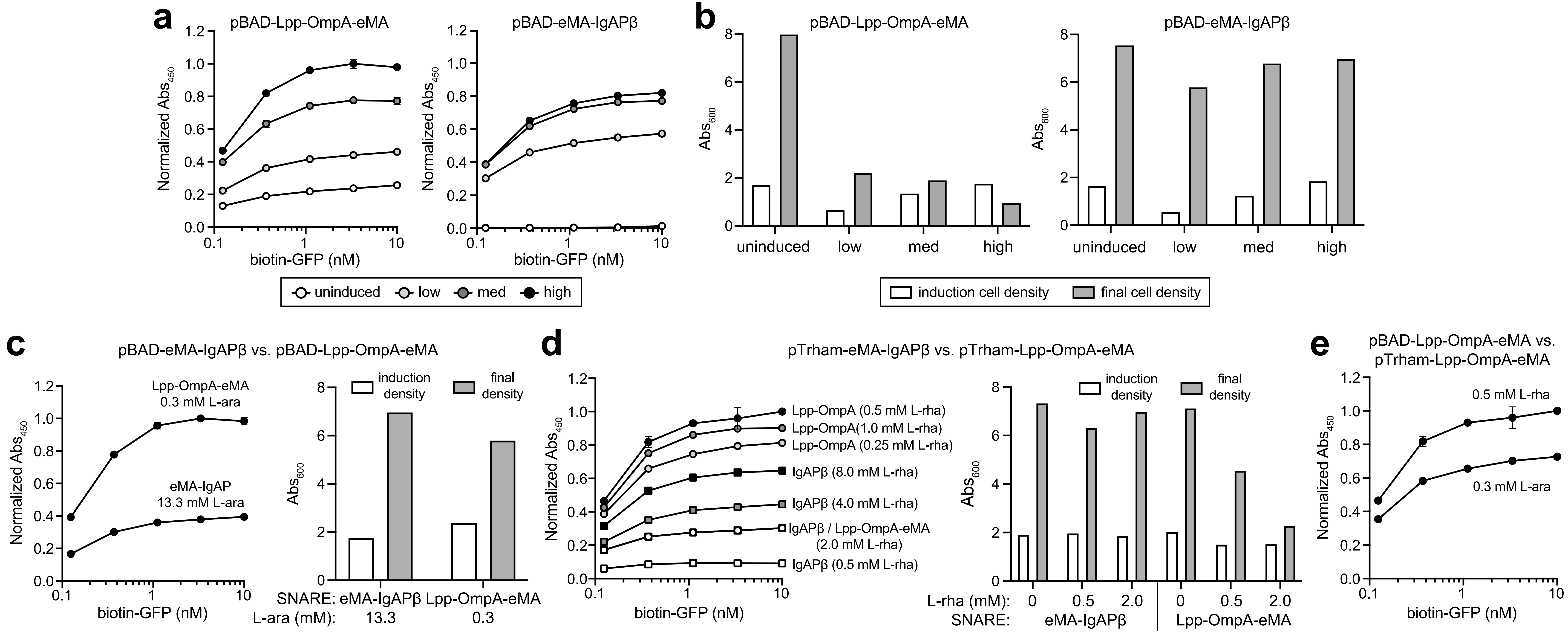


**Supplementary Figure 3. Optimization of biotin-GFP docking on eMA-IgAPβ and Lpp-OmpA-eMA SNAREs.** (a) Binding of biotin-GFP to SNARE-OMVs isolated from hypervesiculating *E. coli* strain KPM404 Δ*nlpI* expressing Lpp-OmpA-eMA or eMA-IgAPβ from plasmid pBAD24 and induced at low ($Abs_{600}$ ~0.6), medium ($Abs_{600}$ ~1.2), or high ($Abs_{600}$ ~1.8) culture density. Data in both graphs were normalized to the maximum binding signal corresponding to Lpp-OmpA-eMA SNARE-OMVs (high induction case) in the presence of 3.3 nM biotin-GFP. (b) Cell growth for same cultures in (a) where cell density was measured at time of induction (white bars) and just prior to harvesting SNARE-OMVs (gray bars). (c) Biotin-GFP binding and cell growth as in (a) and (b) but with 50-fold lower L-arabinose (L-ara) inducer for cells expressing Lpp-OmpA-eMA. Binding data were normalized to the maximum binding signal corresponding to the Lpp-OmpA-eMA SNARE-OMVs in the presence of 3.3 nM biotin-GFP. (d) (left panel) Biotin-GFP binding for SNARE-OMVs isolated from cells expressing Lpp-OmpA-eMA or eMA-IgAPβ from plasmid pTrham in the presence of different amounts of L-rhamnose (l-rha) as indicated. (right panel) Cell growth for a subset of the cells in left panel. (e) Comparison of biotin-GFP binding for Lpp-OmpA-eMA expressed from pBAD24 versus pTrham with inducer amounts as indicated. Data in (d) and (e) were normalized to the maximum binding signal corresponding to the Lpp-OmpA-eMA construct in the presence of 0.5 mM L-rha. Binding activity in all panels was determined by ELISA in which SNARE-OMVs were immobilized on plates and subjected to varying amounts of biotin-GFP, after which plates were extensively washed prior to detection of bound biotin-GFP using anti-polyhistidine antibody to detect C-terminal 6xHis tag on GFP. All binding data are the average of three biological replicates and error bars represent the standard deviation of the mean.

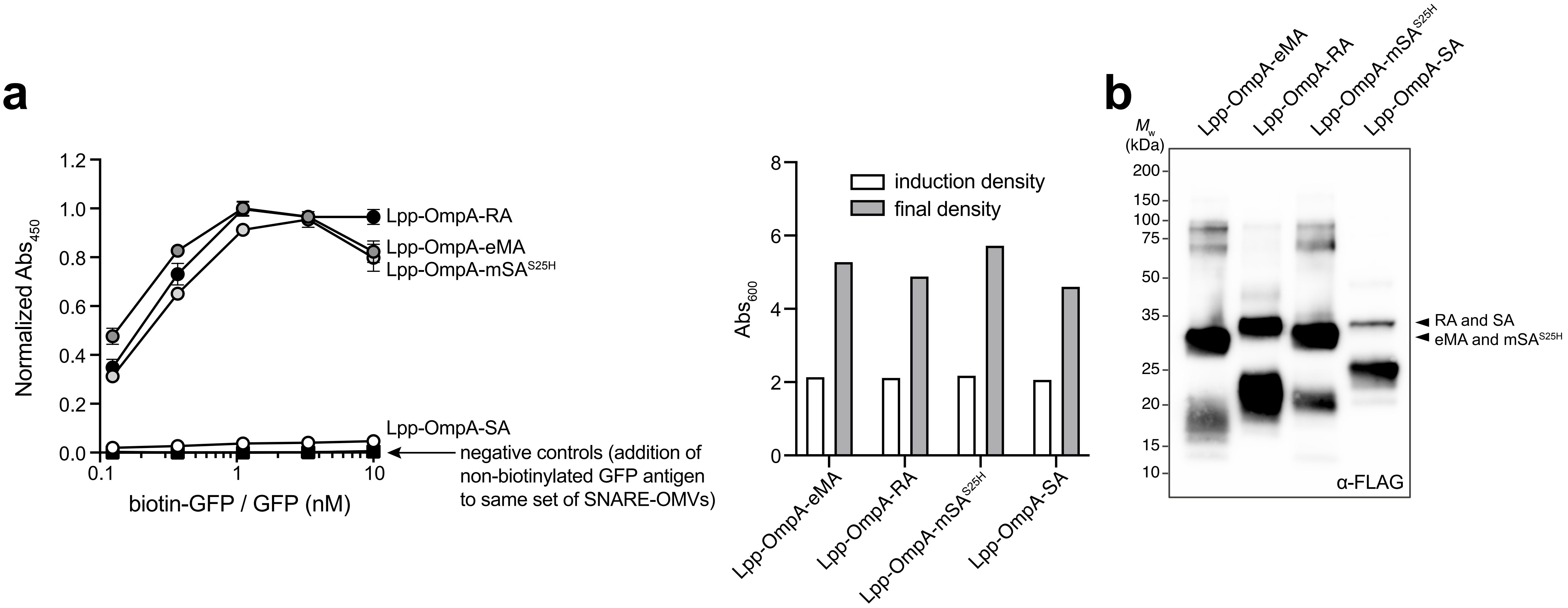


**Supplementary Figure 4. Expression and antigen-binding activity of SNAREs with alternative biotin-binding modules.** (a) (left panel) Binding of biotin-GFP to each of the different SNARE-OMVs as indicated. Binding activity was determined by ELISA in which biotin-binding SNARE-OMVs were immobilized on plates and subjected to varying amounts of biotin-GFP, after which plates were extensively washed prior to detection of bound biotin-GFP using anti-polyhistidine antibody to detect C-terminal 6xHis tag on GFP. Controls were performed by treating the same set of SNARE-OMVs with unmodfied GFP in place of biotin-GFP. All data were normalized to the maximum signal correpsonding to the Lpp-OmpA-eMA construct in the presence of 1 nM biotin-GFP. Datapoints represent the average of three biological replicates and error bars represent the standard deviation of the mean. (right panel) Cell growth for same cultures in (a) where cell density was measured at time of induction (white bars) and just prior to harvesting SNARE-OMVs (gray bars). (b) Immunoblot analysis of OMV fractions isolated from hypervesiculating *E. coli* strain KPM404 Δ*nlpI* expressing each of the different SNAREs from plasmid pBAD24. An equivalent amount of SNARE-OMVs as determined by total protein assay was loaded in each lane. Blot was probed with anti-FLAG antibody (α-FLAG) to detect FLAG epitope (DYKDDDDK) located at the C-terminus of each construct. Expected location of full-length SNARE fusion proteins are denoted by black arrows. Molecular weight ($M_w$) ladder is indicated at left.
